# Cryo-EM structures of human cone visual pigments

**DOI:** 10.1101/2024.01.30.577689

**Authors:** Qi Peng, Jian Li, Haihai Jiang, Xinyu Cheng, Qiuyuan Lu, Sili Zhou, Yuting Zhang, Sijia Lv, Shuangyan Wan, Tingting Yang, Yixiang Chen, Wei Zhang, Weiwei Nan, Tong Che, Yanyan Li, Hongfei Liao, Jin Zhang

## Abstract

Trichromatic color vision in humans constitutes a pivotal evolutionary adaptation, endowing individuals with the capacity to discern and discriminate a diverse spectrum of colors. This unique visual capability confers a selective advantage crucial for successful adaptation, survival, and reproductive success in the natural environment. Color vision in humans is facilitated by the red, green, and blue cone visual pigments within cone photoreceptor cells. These pigments consist of a G-protein-coupled receptor opsin apoprotein and a chromophore covalently linked to opsins. Despite the elucidated structure of rhodopsin, the structures of cone visual pigments have yet to be determined. Here, we present the cryo-EM structures of three human cone visual pigments in complex with G proteins. Our structural analysis reveals detailed interactions between cone opsins, all*-trans*-retinal, and G proteins, indicating their active state. We also provide a concise summary and analysis of mutations in human cone opsins, elucidating potential relationships between residue substitutions and spectral tuning. Notably, S116^2.67^Y, A233^5.52^S, Y277^6.44^F were found to induce a blue shift in the absorption spectrum of the red-pigment, while the substitutions W281^6.48^Y and K312^7.43^A resulted in the absence of the absorption spectrum. The structural elucidation of human cone visual pigments significantly contributes to our understanding of how distinct types of cone cells perceive light across varying wavelengths. Furthermore, it provides a deeper insight into the functioning of the human trichromatic vision system, probing the mechanisms enabling humans to perceive a broad spectrum of colors.

## Introduction

Eye is a most fascinating organ, and the evolution of eye can be traced back to cyanobacteria, the oldest known fossils on the earth. They are the first organisms with ‘eye spots’, organelles for photoreception(Gehring, 2014). Over hundreds of millions of years, the structure of eye has undergone a long and gradual evolution, ultimately giving rise to the sophisticated visual systems in humans(Yokoyama, 2008; Hagen et al., 2023) (Figure 1A). The human eyes rely on the cooperation of cornea and lens to focus light on the retina. Vision in humans, as well as in other vertebrates, is mediated by the rod and cone photoreceptor cells located in the retina of the visual system(Shichida and Matsuyama, 2009; Hofmann and Lamb, 2023) (Figure 1B). In the human retina, there is estimated to be about twenty times more rods (77.9-107.3 million) than cones (4.08-5.29 million)(Curcio et al., 1987; Curcio et al., 1990; Wikler and Rakic, 1990; Srinivasan et al., 2019). The rods mediate scotopic vision working under dim-light conditions, whereas the cones are responsible for photopic vision working daylight conditions(Baylor, 1996; Pichaud et al., 1999; Yokoyama, 2008). In particular, the visual systems in humans and related primates possess a single type of rods, along with three distinct types of cones, thereby enabling trichromatic color vision possible(Kakitani et al., 1985; Nathans et al., 1986; Merbs and Nathans, 1992). This richness in color perception provides humans with a distinct advantage in terms of survival and evolutionary adaption.

**Figure 1:**
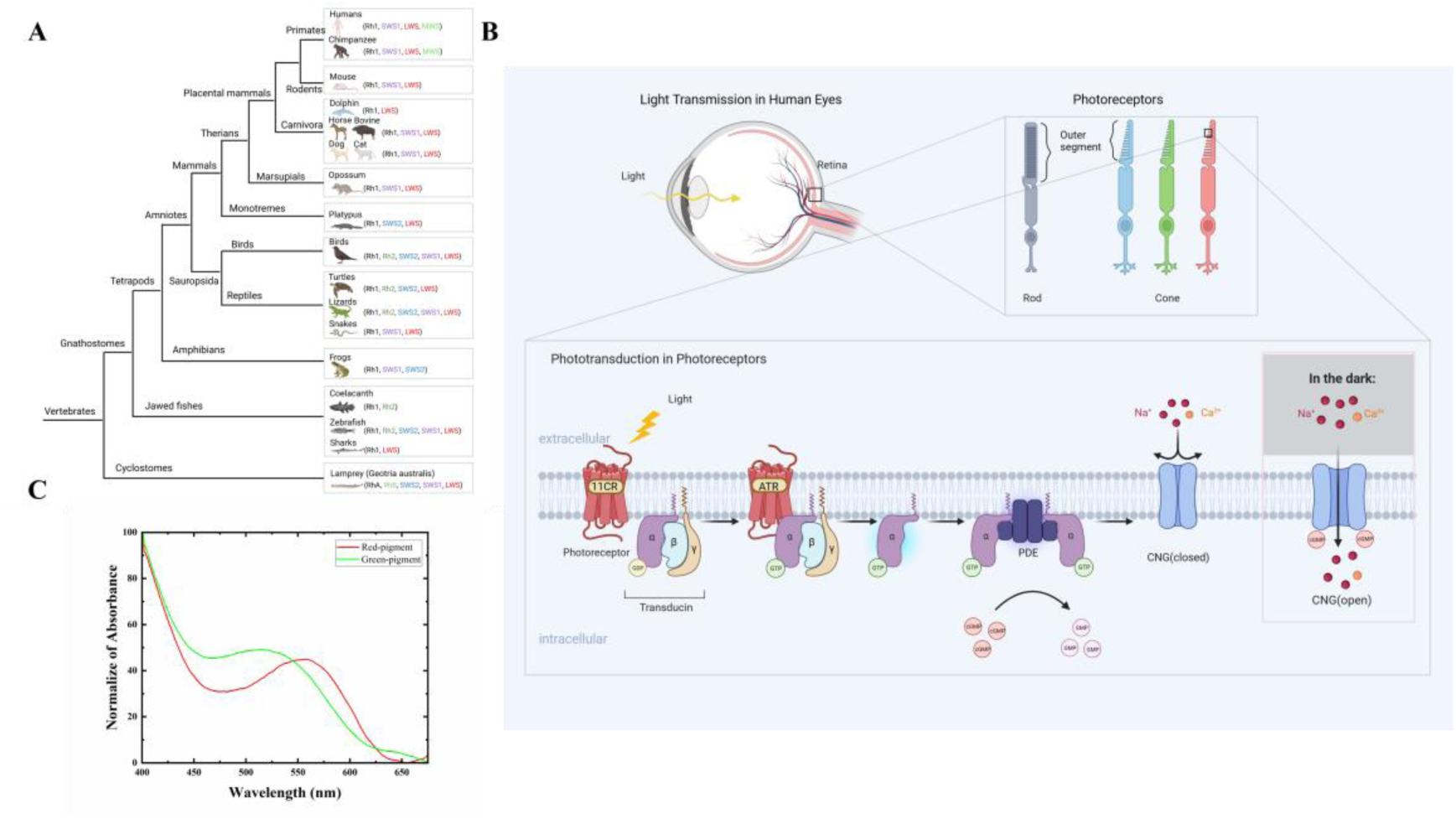
The evolutionary journey of vertebrate photoreceptors and the mechanism of photoreceptor activation. (**A**) The simplified evolutionary tree of photoreceptors in vertebrates, encompassing cyclostomes, gnathostomes, amphibians, birds, reptiles, and mammals, is depicted. The diagram illustrates the various types of photoreceptors present in specific species. Notably, it is crucial to acknowledge that the MWS (medium-wavelength sensitive) photoreceptor discovered in certain primates can be traced back to the ancestral LWS (long-wavelength sensitive) photoreceptor. **(B)** Schematic diagram of the proteins mediating the onset steps of phototransduction, and the flow of activation. When light enters the eye, it is absorbed by 11-*cis* retinal in the visual pigments, which triggers the structural isomerization of the retinal chromophore from 11-*cis* to all-*trans* configuration. This rapid distortion of the chromophore leads to conformational changes in opsins, resulting in their activation (opsins*). Subsequently, the opsins* interacts with the G_t_, which bound to GDP, leading to GDP-GTP exchange and subsequent dissociation of G_t_ into GTP-bound Gα_t_ and Gβγ subunits. The Gα_t_-GTP subunit then interacts with the γ subunit of phosphodiesterase 6 (PDE6), causing conformational changes and activation of PDE6, leading to the hydrolysis of cGMP. As a result, the concentration of cGMP decreases, causing the closure of cyclic nucleotide-gated (CNG) channels. The closure of these channels decreases the influx of cations, hyperpolarizes the cell membrane, and converts the light signal into an electrical signal, which is ultimately transmitted to the brain, resulting in the perception of light. **(C)** UV-visible spectra of red-pigment (red curve) and green-pigment (green curve). The λ_max_ for red-pigment is approximately 560 nm, while the λ_max_ for green-pigment is around 530 nm.

Morphologically, both the rods and cones have an elongated structure and consist of four parts, namely outer segment, inner segment, cell body, and synaptic region. The visual pigments contained in the outer segment is necessary for the absorption of light and subsequent visual sensation(Fain et al., 2010; Lamb, 2022) (Figure 1B). Rods are characterized by the presence of a singular rod visual pigment, rhodopsin. In contrast, cones exhibit a more diverse arrangement with three distinctive cone visual pigments, each featuring a unique absorption maximum. These cone pigments encompass red, green, and blue pigments, composed of long-wave sensitive opsin (LWS-opsin), medium-wave sensitive opsin (MWS-opsin), and short-wave sensitive opsin (SWS-opsin), respectively, each intricately bound to chromophores(Kakitani et al., 1985; Nathans et al., 1986; Merbs and Nathans, 1992). However, wavelength and color are not equivalent. The color we see depends on the relative stimulation of the three cone types. Once there is an abnormality in the function of cones, it will lead to color vision deficiencies.

Color vision deficiencies encompass various types, including red-green defects, such as protanopia (lack of red cones) and protanomaly (replacement of red cones with anomalous pigments), as well as deutan defects arising from the lack of green cones or replacement with anomalous pigments (deuteranomaly). Another class is BCM (Blue Cone Monochromacy), characterized by the loss of both red and green cones. Tritan color vision deficiency, or blue-yellow defect, results from nonfunctional blue cones. Finally, achromatopsia, or complete color blindness, is attributed to the loss of function in all three classes of cones. As color discrimination predominantly depends on opsins in cones, elucidating the structure of human cone opsins contributes to a better understanding of vision-related diseases. For instance, mutations in cone opsins may be associated with certain hereditary visual impairments(Neitz et al., 2020; Zhu et al., 2021), and structural studies will assist in revealing how these mutations affect visual function.

Visual pigments found in photoreceptor cells is composed of opsin and 11-*cis* retinal (a derivative of vitamin A), which are covalently linked together(Nathans et al., 1986; Kosower, 1988; Zhukovsky and Oprian, 1989). For most vertebrates, the chromophore, 11-*cis* retinal, in rod and cone visual pigments is the same. Despite containing the same 11-*cis*-retinal chromophore, differences exist in the structures of rod and cone opsin apoproteins, resulting in the unique spectral response of photoreceptor cell type(Imamoto and Shichida, 2014). The opsin is a type of G-protein-coupled receptor (GPCR) with seven transmembrane helical structures(Sakmar and Khorana, 1988; Terakita, 2005; Palczewski, 2006; Ernst et al., 2014). When light enters the eye, it is absorbed by 11-*cis* retinal in the visual pigments, which triggers the structural isomerization of the retinal chromophore from 11-*cis* to all-*trans* configuration. This rapid distortion of the chromophore leads to conformational changes in opsins, resulting in their activation (opsins*)(Hubbard and Kropf, 1958). The activated opsins* then interact with GDP-bound transducin (a heterotrimeric G protein known as G_t_), leading to GDP-GTP exchange and subsequent dissociation of G_t_ into GTP-bound Gα_t_ and Gβγ subunits. The Gα_t_-GTP subunits then interact with the γ subunits of phosphodiesterase 6 (PDE6), causing conformational changes and activation of PDE6. This activation leads to the hydrolysis of cGMP(Miki et al., 1975; Hubbell and Bownds, 1979; Liebman and Pugh, 1979; Cote, 2021). As a result, the concentration of cGMP decreases, causing the cyclic nucleotide-gated (CNG) channels to close(Caretta et al., 1985; Fesenko et al., 1985; Koch and Kaupp, 1985). The closure of these channels reduces the influx of cations, hyperpolarizes the cell membrane, and converts light signals into electrical signals that are ultimately transmitted to the brain, resulting in the perception of light(Barret et al., 2022; Chen et al., 2022) (Figure 1B).

Previously, the structure of bovine rhodopsin was solved at the atomic level using X-ray crystallography, which is the first solved GPCR structure(Palczewski et al., 2000). A wealth of structural information of both human and bovine rhodopsin is currently available, including the ligand-free state, inactive state, and active state structures(Palczewski et al., 2000; Okada et al., 2004; Park et al., 2008; Choe et al., 2011; Zhou et al., 2017; Kang et al., 2018; Tsai et al., 2018; Tsai et al., 2019; Gao et al., 2019; Chen et al., 2021). These data are of great value for the understanding of phototransduction. However, there is a shortage of research on the human cone opsins, and the structures of cone opsins have not been elucidated yet.

In this study, we determined the cryo-EM structures of human LWS-, MWS-, and SWS-opsins in complex with all-*trans* retinal and G_i_ proteins. Detailed structural analysis and comparison revealed the interaction between cone opsins and G proteins as well as their chromophore, which provides a solid foundation to understand molecular mechanisms behind human perception of different colors. Moreover, the present study summarized and analyzed the residue substitution of human cone opsins, which indicated that certain mutation of single amino acid could cause a spectral shift in wavelength sensitivity and relate with human color vision defects.

## Results

### Structural determination of cone opsin-G_i_ complexes

To determine the structure of cone opsins, the agonist all-*trans* retinal was used to form LWS-opsin-G protein, MWS-opsin-G protein and SWS-opsin-G protein complexes. By aligning the sequence with rhodopsin, we introduced a constitutive active mutation E129^3.28^Q [superscript indicates Ballesteros-Weinstein nomenclature] into the full-length wild-type OPN1LW/OPN1MW (constitutive active mutation E110^3.28^Q in OPN1SW)(Zhukovsky and Oprian, 1989; Sakmar et al., 1989; Robinson et al., 1992; Standfuss et al., 2011) (Figures S1 and S2A). Previous researches have shown that the active rhodopsin-G_i_ complex is more stable than the active rhodopsin-G_t_ complex(Maeda et al., 2014), so we focused on obtaining a stable complex of active LWS-opsin/MWS-opsin/SWS-opsin with a dominant-negative form of G_i_ (DNGα_i_), which was used to decrease the affinity of nucleotide-binding(Liu et al., 2016; Draper-Joyce et al., 2018) (Figure S2). The NanoBiT tethering strategy(Duan et al., 2020; Zhou et al., 2020) was applied to stabilize the LWS-opsin-G_i_/MWS-opsin-G_i_/SWS-opsin-G_i_ complex, which has been widely used in the structural determination of GPCR-G protein complexes. Specifically, the C-terminus of the receptor was linked to the large subunit LgBiT, while the C-terminus of the human Gβ subunit was connected to the small subunit HiBiT. In addition, scFv16 was used to further stabilize the complexes, according to previous studies(Koehl et al., 2018; Mao et al., 2020). The LWS-opsin-G_i_/MWS-opsin-G_i_/SWS-opsin-G_i_ complexes were assembled by co-expressing the engineered receptor with DNGα_i1_, Gβ_1_γ_2_ subunits, and scFv16 in Sf9 cells. During the process of purification, the agonist all-*trans* retinal was added, yielding the complex protein. Meanwhile, we individually purified the LWS-opsin and MWS-opsin with the addition of 11-*cis* retinal, with absorption wavelengths of 560 nm and 530 nm, respectively (Figure 1C). These absorption spectra closely resemble the previously reported data(Kakitani et al., 1985; Nathans et al., 1986; Merbs and Nathans, 1992), providing evidence that the cone pigments reconstituted in vitro exhibit physiological properties similar to their cellular counterparts.

Cryo-EM analysis with extensive particle classification yielded high-resolution three-dimensional (3D) density maps for LWS-opsin-G_i_, MWS-opsin-G_i_, and SWS-opsin-G_i_, with nominal global resolutions of 3.4 Å, 2.5 Å, and 2.6 Å, respectively (Figure 2, Figures S4 to S6, Table S1). The high-quality density maps are clear for modeling LWS-opsin/MWS-opsin from residue 38 to residue 335 and SWS-opsin from residue 37 to residue 315. Other amino acid residues at the N-terminus and C-terminus of the receptors are too flexible and disordered, making them unable to be used for modeling. In contrast, all-*trans* retinal, scFv16 and the three subunits of G_i_ protein are well-fitted in the EM map.

**Figure 2:**
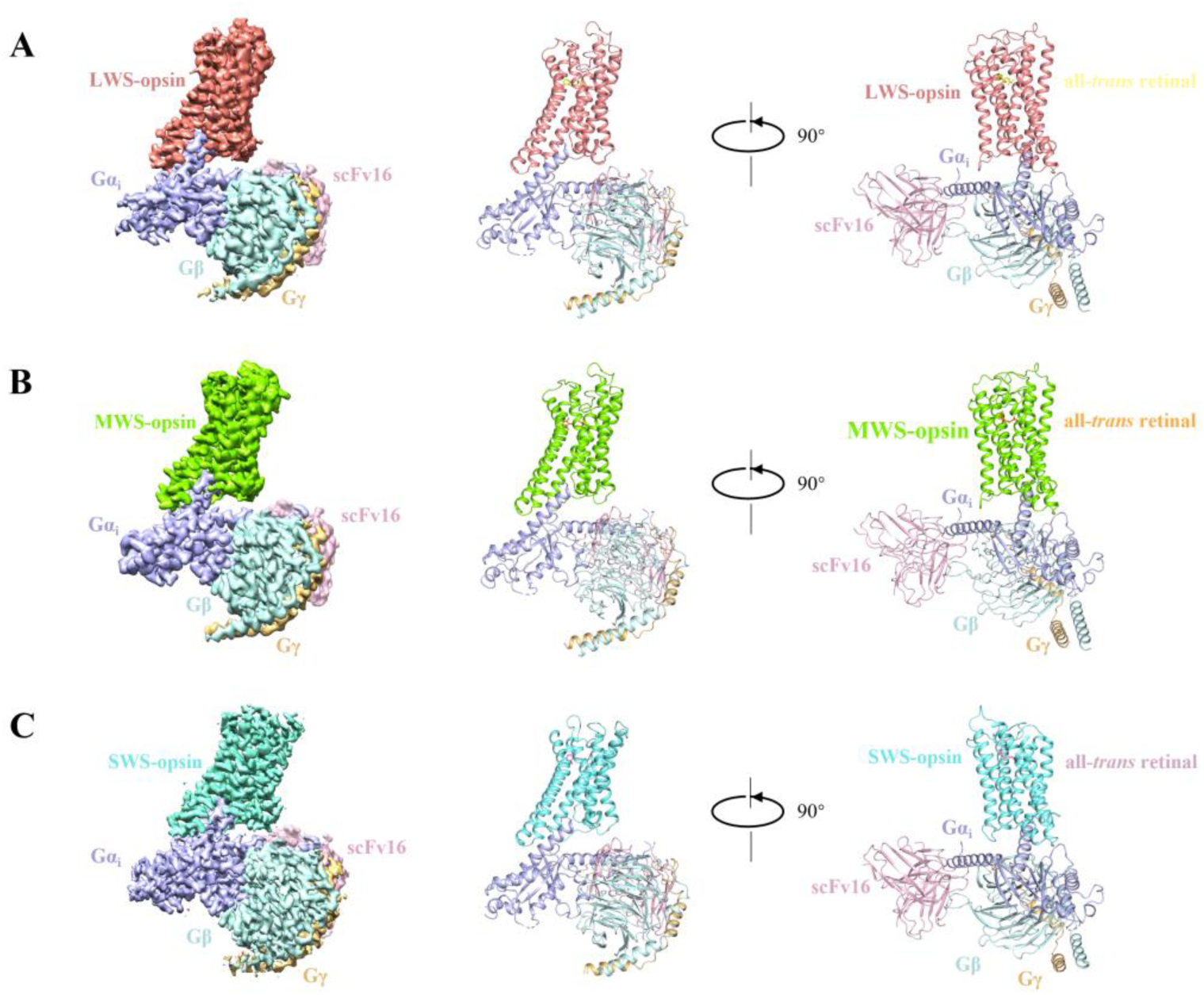
Overall architecture of LWS-opsin-G_i_, MWS-opsin-G_i_ and SWS-opsin-G_i_. (**A**) Cryo-EM map and cartoon model of the LWS-opsin-G_i_ complex colored by subunit (LWS-opsin in salmon, Gα_i_ in light blue, Gβ_1_ in pale cyan, Gγ_2_ in light orange, scFv16 in light pink, all-*trans* retinal in tv-yellow). **(B)** Cryo-EM map and cartoon model of the MWS-opsin-G_i_ complex colored by subunit (MWS-opsin in chartreuse, all-*trans* retinal in tv-orange). **(C)** Cryo-EM map and cartoon model of the SWS-opsin-G_i_ complex (SWS-opsin in aquamarine, all-*trans* retinal in pink). The subunits of G protein and scFv16 are indicated with the same color as in the LWS-opsin-G_i_.

Globally, the structures of cone opsin-G_i_ are similar to the previously reported rhodopsin-G_i_/G_t_ protein structures, but provide more detailed structural information of distinct visual pigments. These findings establish a solid structural foundation for advancing our understanding of the molecular mechanisms underlying the perception of different colors.

### Retinal binding pocket

The complex structures provide insight into retinal-cone opsins interactions in the ligand binding site. In our structures, all-*trans* retinal occupies the orthosteric binding site. The polyene chain of all-*trans* retinal extends towards the residue K312^7.43^ in LWS-opsin/MWS-opsin (K293^7.43^ in SWS-opsin), allowing the carbon atom C15 of retinal to form a Schiff base with the ε-amino group of the conserved lysine at position 7.43 on TM7 (Figure 3). This interaction is supported by the strong density connecting retinal and the side chain of K^7.43^ (Figures 3A, 3D and 3G), along with our UV-visible spectroscopy data (Figure S3). In all three structures, the residues involved in the formation of the ligand-binding pockets are primarily located in TM2, TM3, TM5, TM6, TM7, and ECL2 (Figure 3).

**Figure 3:**
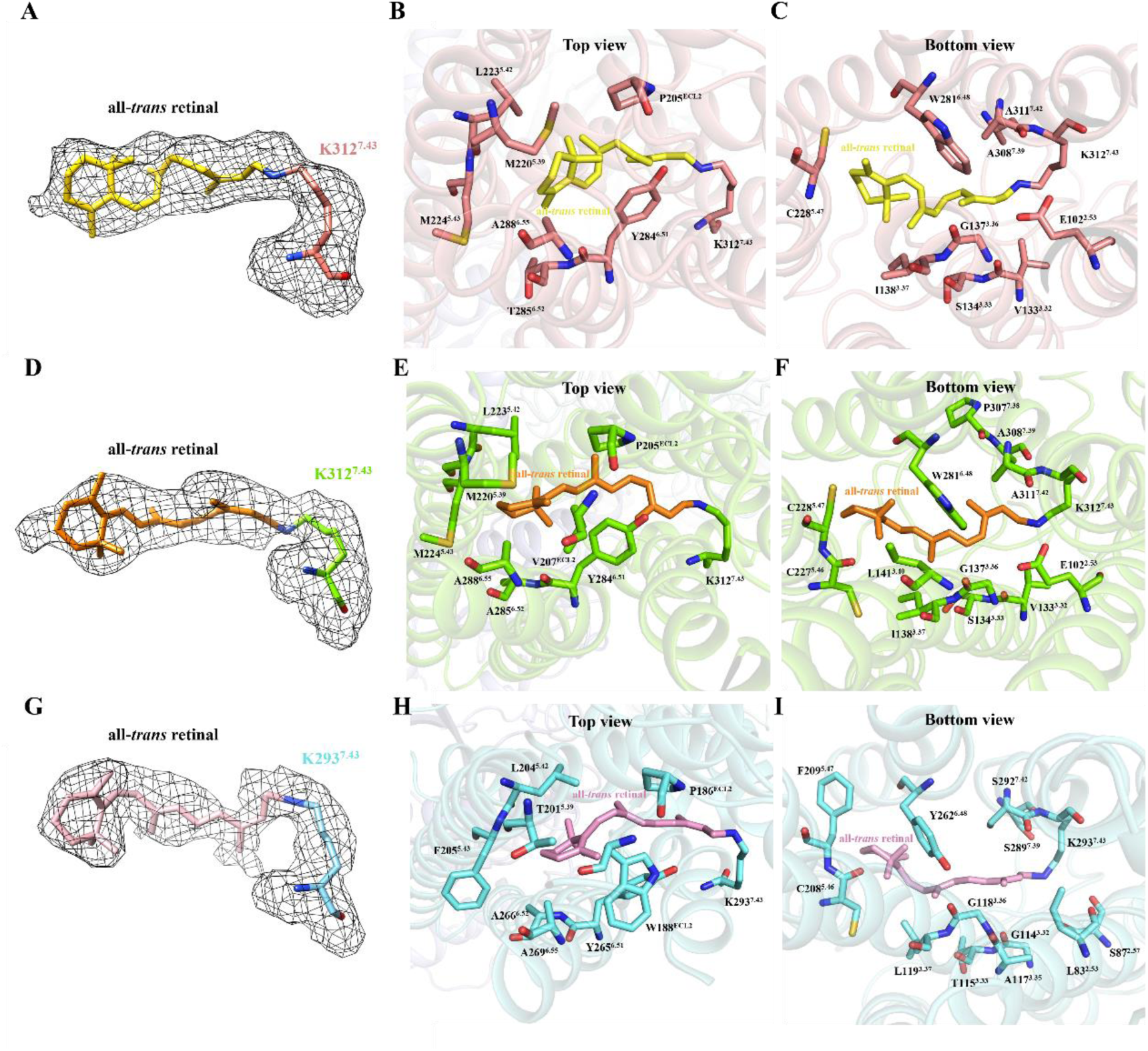
Retinal binding pocket of LWS-opsin-G_i_, MWS-opsin-G_i_ and SWS-opsin-G_i_. (**A**) The density of all-*trans* retinal in the LWS-opsin-G_i_ is contoured at 2.0 σ. **(B-C)** The retinal binding pocket of LWS-opsin-G_i_ is illustrated from both the top **(B)** and bottom views **(C)**, featuring LWS-opsin in salmon and all-*trans* retinal in tv-yellow.**(D)** The density of all-*trans* retinal in the MWS-opsin-G_i_ is contoured at 2.0 σ. **(E-F)** The retinal binding pocket of MWS-opsin-G_i_ is presented from both the top **(E)** and bottom views **(F)**, showcasing MWS-opsin in chartreuse and all-*trans* retinal in tv-orange.**(G)** The density of all-*trans* retinal in the SWS-opsin-G_i_ is contoured at 2.0 σ. **(H-I)** The retinal binding pocket of the SWS-opsin structure is also showcased from the top view **(H)** and bottom view **(I)**, highlighting SWS-opsin in aquamarine and all-*trans* retinal in pink.

Comparing the residues in the ligand-binding pocket of LWS-opsin with MWS-opsin, we observed a difference only at residue 285^6.52^. Specifically, LWS-opsin has a T285^6.52^, while MWS-opsin has an A285^6.52^ (Figures 3B and 3E). According to previous researches, this amino acid is one of the key residues contributing to the major difference of the absorption maximum(Yokoyama and Yokoyama, 1990; Neitz et al., 1991). It has been revealed that in Rhodopsin, the counterion of K296^7.43^/Schiff base is E113^3.28^ in the resting state and E181^ECL2^ in the metarhodopsin conformation.. This conservation extends to SWS-opsin, where the corresponding residues are E110^3.28^ and E178^ECL2^ (Figure S1). However, LWS-opsin and MWS-opsin differ from SWS-opsin in lacking E^ECL2^ and have a E102^2.53^ instead, which is in proximity to K312^7.43^ and E129^3.28^. As a result, the potential ionic interactions can be formed between K^7.43^/Schiff base and E^3.28^ or E^ECL2^ in rhodopsin and SWS-opsin, while E^3.28^ or E^2.53^ in LWS-opsin and MWS-opsin. In our study, we confirmed the spatial position of E102^2.53^ in relation to K312^7.43^ in the LWS-opsin and MWS-opsin structures (Figures 3C and 3F).

Previous studies have shown that the simultaneous substitution of nine specific amino acids (M86^2.53^L, G90^2.57^S, A117^3.32^G, E122^3.37^L, A124^3.39^T, W265^6.48^Y, A292^7.39^S, A295^7.42^S, and A299^7.46^C) in rhodopsin resulted in a significant blue-shift in the absorption maximum(Lin et al., 1998). In our SWS-opsin-G_i_ structure, it can be observed that some of the residues (L83^2.53^, S87^2.57^, G114^3.32^, L119^3.37^, Y262^6.48^, S289^7.39^, and S292^7.42^) are located within the ligand-binding pocket (Figures 3H and 3I). In particular, the presence of non-hydroxyl-bearing amino acids or hydroxyl-bearing amino acids, such as Serine or Threonine, often affects spectral sensitivity. For instance, when a hydroxyl-bearing amino acid is located near the β-ionone ring of the retinal, it helps to maintain the excited state of the chromophore. This results in a lower energy requirement for isomerization, leading to opsins that predominantly absorb long-wavelength/low-energy light, commonly found in the red spectrum. On the other hand, the presence of S/T near the Schiff-base site stabilizes the ground state of the retinal, requiring the activation of short-wavelength/high-energy light, typically found in the blue spectrum(Hagen et al., 2023). Our structure reveals that residues S87^2.57^, S289^7.39^ and S292^7.42^ are located in proximity to the Schiff-base region, while T201^5.39^ is positioned near the β-ionone ring of the retinal (Figures 3H and 3I). A conserved motif, CW^6.48^xP, exists in GPCRs, with W^6.48^ serving as the “toggle switch” during GPCR activation. However, the corresponding residue in SWS-opsin is Y262^6.48^. (Figures 3I and 4C).

**Figure 4:**
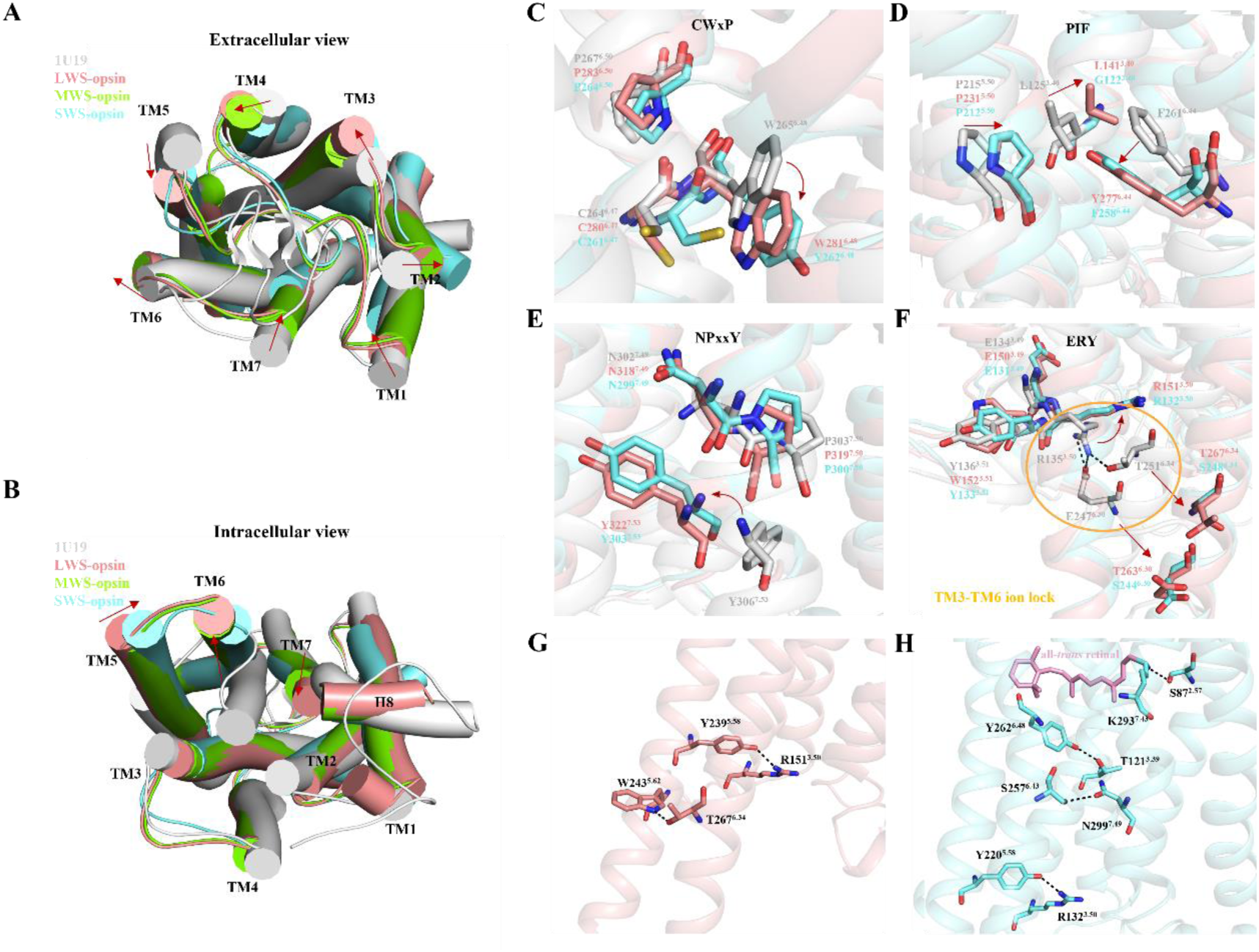
Structural comparison of cone opsins and inactive rhodopsin. (**A-F**) Comparison of cone opsins in our structures with inactive Rhodopsin (PDB ID 1U19), including the overall structure of transmembrane bundle from the extracellular view **(A)** and intracellular view **(B)**, as well as close-up views of the conformational changes of “toggle switch”-W^6.48^ **(C)**, rearrangement of “PIF” motif **(D)**, rearrangement of Y^7.53^ in “NPxxY” motif **(E)**, and conformational changes of “ERY” motif **(F)**. **(G-H)** Polar contacts in LWS-opsin **(G)** and SWS-opsin **(H)** are shown as black dashed lines. LWS-opsin, MWS-opsin, SWS-opsin and inactive Rhodopsin are shown in salmon, chartreuse, aquamarine and gray respectively. The red arrows indicate conformational changes of conserved motifs between active cone opsins and inactive Rhodopsin.

### Activation of cone opsins

In our structural analysis, a notable resemblance is evident between the LWS-opsin-G_i_ and MWS-opsin-G_i_ structures, with a root mean square deviation (RMSD) of 0.609 Å for equivalent Cα positions. Meanwhile, the RMSD value between the LWS-opsin-G_i_ and SWS-opsin-G_i_ structures is 0.975 Å. Comparing our structures with the inactive rhodopsin (PDB ID 1U19), Several conformational differences have been identified. These include an inward movement of TM1, 5, and 7, coupled with an outward movement of TM 2, 3, and 6 when viewed from the extracellular side (Figure 4A). The most notable differences, however, involve the elongation of TM5 and the outward shift of TM6 on the cytoplasmic side (Figure 4B). In the LWS-opsin-G_i_ and MWS-opsin-G_i_ structures, TM6 undergoes an outward movement of approximately 11 Å, demonstrating similar displacement. Similarly, the SWS-opsin-G_i_ structure exhibits an approximately 10 Å outward shift in TM6 (Figure 4B). Given the substantial similarity between LWS-opsin and MWS-opsin structures, we conducted a structural analysis of LWS-opsin and SWS-opsin, with a particular focus on the conserved motifs in GPCRs (Figure 4). It was revealed that isomerization of 11-*cis* retinal to all-*trans* retinal occurs, resulting in the displacement of β-ionone from residue W281^6.48^/Y262^6.48^. This displacement reduces the packing between the β-ionone and W281^6.48^/Y262^6.48^, inducing a flip of this residue (Figure 4C). The residue W^6.48^, known as the “toggle switch”, initiates a rearrangement of the PIF motif, leading to a deviation of Y^7.53^ within the NPxxY motif (Figures 4D and 4E). In inactive rhodopsin, there are two pairs of interactions, R135^3.50^-E247^6.30^ and R135^3.50^-T251^6.34^, forming the TM3-TM6 ionic lock. However, as a result of receptor activation and the outward movement of TM6, the ionic lock is disrupted (Figure 4F). Consequently, in our cone opsin structures, R151^3.50^/R132^3.50^ has been liberated from the ionic lock. On the other hand, TM5 undergoes extension, Y239^5.58^/Y220^5.58^ undergoes a significant rotational shift towards TM3, which exposes its side chain to the nearby environment of R151^3.50^/R132^3.50^, and thus allows it to serve as a hydrogen bonding receptor for the guanidine moiety of R151^3.50^/R132^3.50^. Consequently, R151^3.50^/R132^3.50^ and Y239^5.58^/Y220^5.58^ have formed a structurally significant interaction to stabilize the active state of the receptor (Figures 4G and 4H). Furthermore, the hydrogen bond between W243^5.62^ and T267^6.34^ in the LWS-opsin-G_i_ structure solidifies the linkage between TM5 and TM6 (Figure 4G), bringing the two transmembrane helices into closer contact with one another. Overall, the conformation of SWS-opsin-G_i_ is similar to that of LWS-opsin-G_i_ (Figure 4). In more detail, there is elongation of TM5, outward movement of TM6, leading to the disruption of the TM3-TM6 ionic lock, and Y220^5.58^ creating a hydrogen bond with R132^3.50^ (Figure 4H). However, there are still some structural disparities. In SWS-opsin, S87^2.57^ forms a hydrogen bond with the conserved K293^7.43^ in the opsin family. Additionally, S257^6.43^ interacts with N299^7.49^ of the NPxxY motif (Figure 4H). It has been known that T121^3.39^ is involved in spectral tuning, but in our structure, it is not located within the ligand binding pocket, but there is an interplay between T121^3.39^ and Y262^6.48^ (Figure 4H), therefore, we speculate that it may participate in spectral tuning by interacting with Y262^6.48^.

### Interface between cone opsins and G_i_

By superimposing our cone opsin-G_i_ structures onto rhodopsin-Gt complex (PDB ID 6OYA), an undeniable resemblance in the overall architecture becomes apparent (Figure 5A), with the RMSD value of equivalent Cα positions for LWS-opsin-G_i_, MWS-opsin-G_i_, and SWS-opsin-G_i_ being 1.235 Å, 1.156 Å, 1.183 Å, respectively. When comparing the structure of LWS-opsin-G_i_ with that of the rhodopsin-Gt complex, the α5 helix inserts consistently into the receptor depth. However, the αN helix in LWS-opsin-G_i_ undergoes an upward shift of 2.2 Å (Figure 5B). Our structural analysis highlights key regions in G_i_ binding to receptors, encompassing the αN helix, α4-β6 loop, β6 sheets, and α5 helix (Figure S7). The main contact regions in receptors are situated at the cytoplasmic end of TM2, TM3, TM5, TM6, and TM7/H8 joint, as well as the ICL2 and ICL3 (Figure S7). In these structures, the primary interface between G_i_ and the cone opsin is mediated by the C-terminal α5 helix of Gα_i_. The residues 331-354 of Gα_i_ adopt a straight amphipathic helix that inserts into the open cytoplasmic end of the receptors, with the final four residues forming a “hook”. In the LWS-opsin-G_i_ complex, several pairs of polar interactions are evident, including Q253^ICL3^-Y320^h4s6.12^/D341^H5.11^, S258^6.25^-K345^H5.17^, R327^8.48^-F354^H5.26^, and Q328^8.49^-K349^H5.21^/D350^H5.22^.

**Figure 5:**
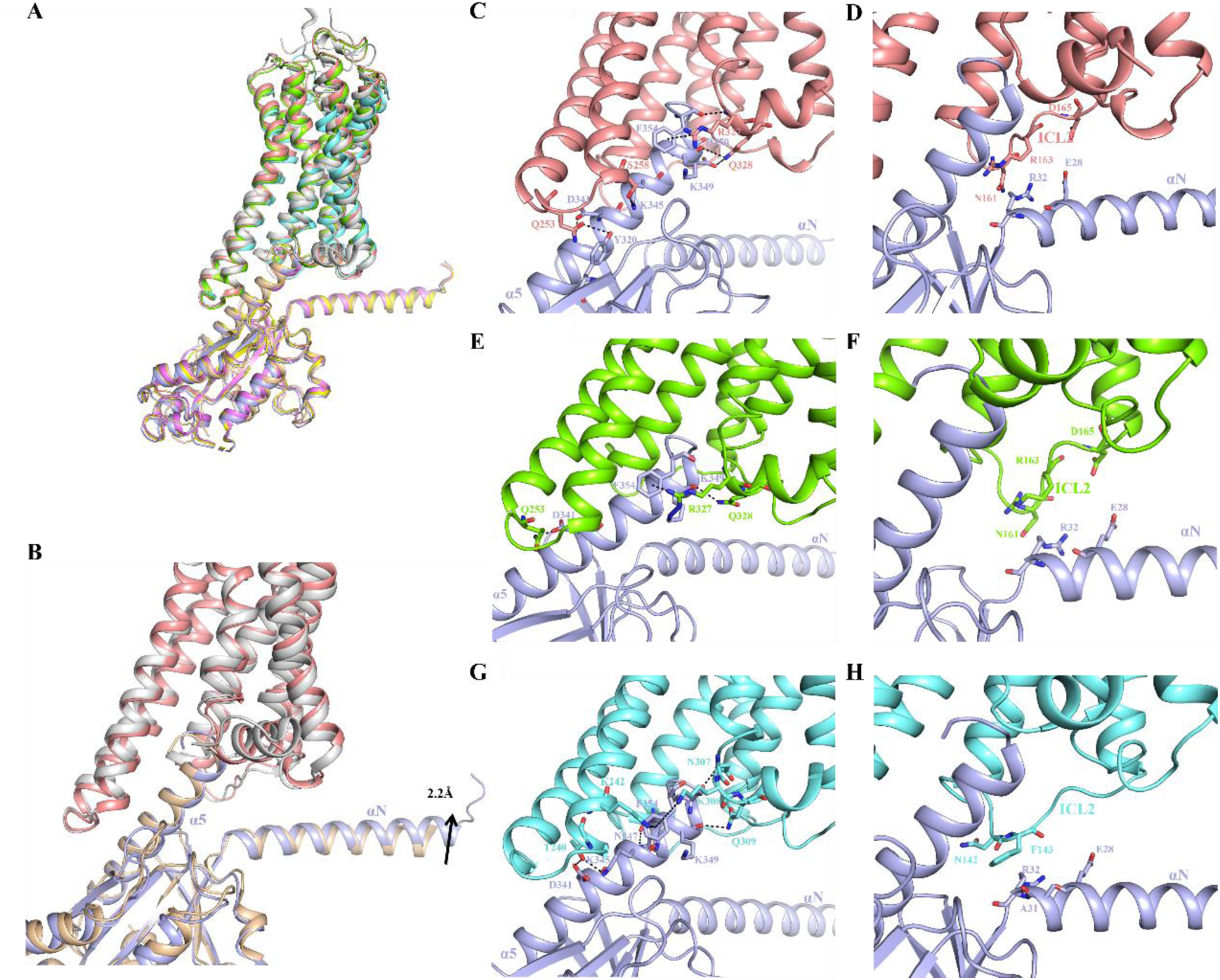
Interface between cone opsins and G_i_. (**A**) Structural comparison of the LWS-opsin-G_i_, MWS-opsin-G_i_, SWS-opsin-G_i_ and Rhodopsin-G_t_ (PDB ID 6OYA), with Gβγ and antibodies omitted. The LWS-opsin, MWS-opsin, SWS-opsin, and Rhodopsin are colored by salmon, chartreuse, aquamarine, and gray respectively, while the Gα subunits are represented by light blue, yellow, violet, and wheat separately. **(B)** Close-up view of superposition of the LWS-opsin-G_i_ and Rhodopsin-G_t_ (PDB ID 6OYA) with LWS-opsin represented in salmon, G_i_ in light blue, Rhodopsin in gray, and G_t_ in wheat. **(C-D)** The primary interface **(C)** and the secondary interface **(D)** between Gα_i_ and LWS-opsin. **(E-F)** The primary interface **(E)** and the secondary interface **(F)** between Gα_i_ and MWS-opsin. **(G-H)** The primary interface **(G)** and the secondary interface **(H)** between Gα_i_ and SWS-opsin.

Additionally, a π-cation interaction is observed between the residue F354^H5.26^ in Gα_i_ and the positively charged amino acid residue R327^8.48^ in LWS-opsin (Figure 5C). Similarly, in the MWS-opsin-G_i_ structure, there are two pairs of hydrogen bond interactions, Q253^ICL3^-D341^H5.11^ and Q328^8.49^-K349^H5.21^, and a π-cation interaction between F354^H5.26^ and R327^8.48^ (Figure 5E). As for SWS-opsin-Gi, polar interactions (V135^3.53^-N347^H5.19^, T240^6.26^-D341^H5.11^/K345^H5.15^, N307^8.47^-C351^H5.23^ and Q309^8.49^-K349^H5.21^), along with π-cation interactions (F354^H5.26^-K242^6.28^/K308^8.48^), have also been identified (Figure 5G). In addition, we identified the secondary interface between G_i_ and cone opsins, which is formed by αN helix of Gα_i_ and the ICL2 of cone opsins. Specifically, the residues E28 and R32 of the αN helix interact with ICL2 (in SWS-opsin-G_i_, A31 is also involved in forming the secondary interface) (Figures 5D, 5F and 5H). It is important to note that this interaction between E28 and R32 and the receptor’s ICL2 was also observed in the rhodopsin-G_i_ structure(Kang et al., 2018).

### Implication for color vision deficiencies

Mutations of the genes encoding visual pigments may cause the insufficient function of the visual systems and result in color vision deficiencies such as red-green color blindness, which is the most common type of color vision deficiencies. In this study, we collected some mutations associated with color vision deficiencies (Figure 6 and Figure S8). The mutations identified within the LWS-opsin are K82^ICL1^E, N94^2.45^K, V120^ECL1^M, S143^3.42^P, W177^4.50^R, C203^ECL2^R, P231^5.50^L, R247^5.66^X, P307^7.38^L, R330^8.51^Q, and G338^8.59^E(Nathans et al., 1993; Ueyama et al., 2002; Carroll et al., 2004; Ueyama et al., 2004; Mizrahi-Meissonnier et al., 2010; Cai et al., 2019; Iarossi et al., 2021; Zhu et al., 2021) (Figures 6A and S8A). It is noteworthy that the C203^ECL2^R mutation disrupts the disulfide bond linkage between C203^ECL2^ and C126^3.25^, which is also observed in MWS-opsin (Figure 6B). Furthermore, the residues W177^4.50^, N94^2.45^, and S143^3.42^ collaboratively form a hydrogen bonding network (Figure 6A), contributing to the stabilization of the receptor conformation. Mutations of these residues may reduce protein stability, subsequently leading to associated diseases. Moreover, the R247^5.66^X substitution introduces a stop codon, leading to aberrant protein translation and consequently causing color vision deficiencies. For MWS-opsin, in accordance with LWS-opsin, the mutation C203^ECL2^R disrupts the disulfide bond between C203^ECL2^ and C126^3.25^ (Figure 6B). Moreover, N94^2.45^K, W177^4.50^R, P307^7.38^L, R330^8.51^Q, and G338^8.59^E mutations can also lead to color vision deficiency(Ueyama et al., 2002; Zhu et al., 2021) (Figures 6B and S8B). The MWS-opsin missense mutation P187^4.60^S has been identified in subjects with color-vision deficiency(Neitz et al., 2004), and P187^ECL2^ is conserved among vertebrate visual pigments (Figure S1). Replacing this conserved proline residue with serine is likely to disrupt the proper folding and function of the photopigment molecule. For SWS-opsin, six mutations have been identified as associated with tritan color vision deficiency: L56^1.54^P, G79^2.49^R, T190^ECL2^I, S214^5.52^P, P264^6.50^S, and R283^7.33^Q(Weitz et al., 1992; Neitz et al., 2020). These mutations may lead to significant steric hindrance and disrupt the stability of the SWS-opsin (Figures 6C and S8C), and the mutation P264^6.50^S potentially influences the maximum absorption wavelength of the pigment by affecting Y262^6.48^.

**Figure 6:**
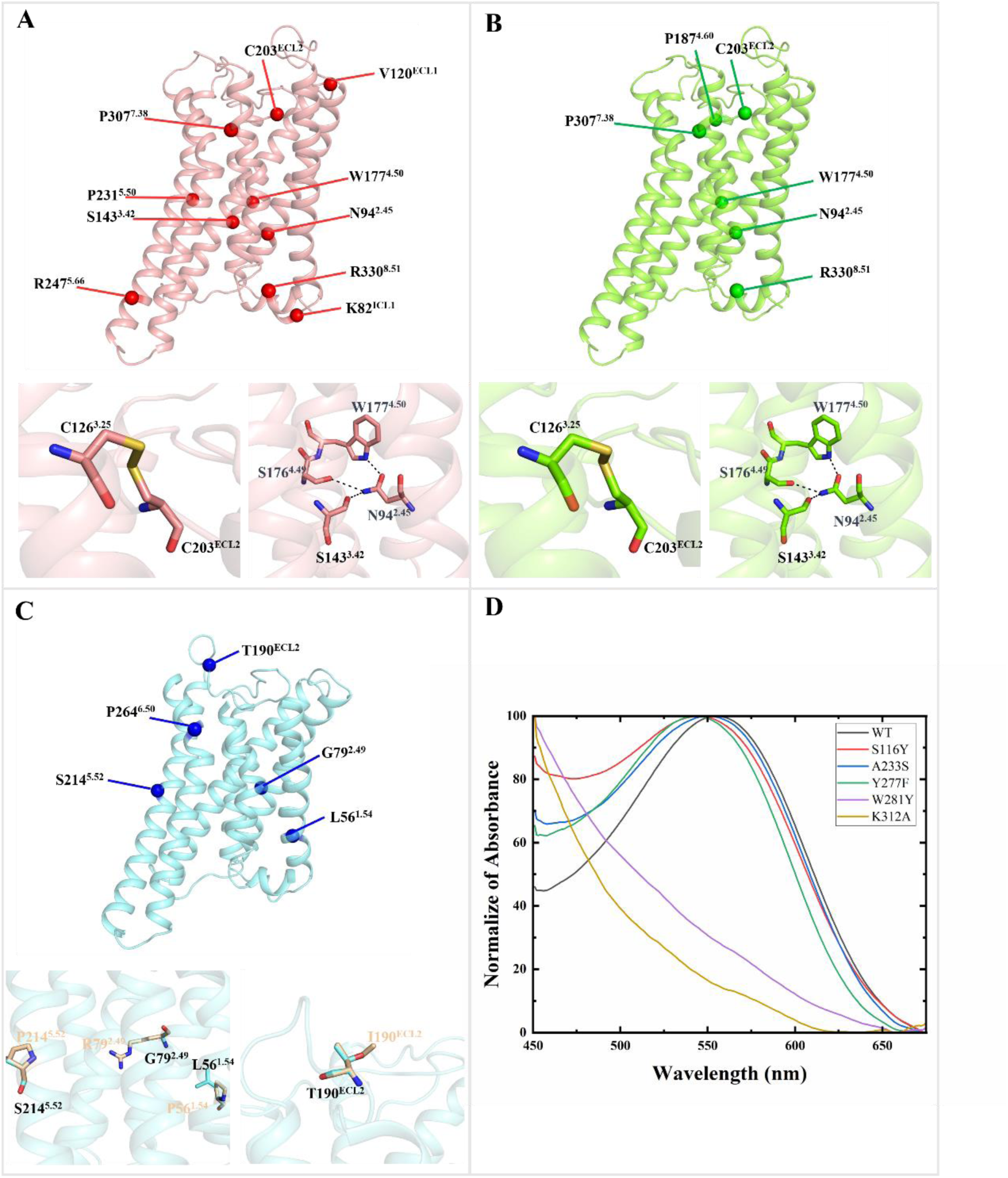
Color Vision Deficiency Mutations in Cone Opsins. Color vision deficiency-related mutations associated with cone opsins are mapped onto the structure of cone opsins. **(A)** Mutations in LWS-opsin linked to color vision defects are represented by red spheres. **(B)** Mutations in MWS-opsin associated with color vision defects are denoted by green spheres. **(C)** Mutations in SWS-opsin related to color vision defects are indicated by blue spheres, with residues before and after the mutation displayed in chartreuse and wheat, respectively. LWS-opsin, MWS-opsin, and SWS-opsin are represented by salmon, chartreuse, and aquamarine ribbons, respectively and polar contacts are shown as black dashed lines. **(D)** UV-visible absorption spectra of red-pigment (WT and mutants). Residues in LWS-opsin were substituted with the corresponding residues from MWS-opsin (S116^2.67^Y, A233^5.52^S, Y277^6.44^F), inducing a blue shift of λ_max_, bringing it closer to that of MWS-opsin. However, W281^6.48^Y and K312^7.43^A resulted in the absence of absorption wavelengths resembling those of either red-pigment or green-pigment.

In order to gain a more in-depth understanding of the specific amino acid residues crucial for influencing the absorption spectra of cone-opsins, systematic mutagenesis experiments were conducted, involving the substitution of LWS-opsin residues with their MWS-opsin counterparts based on the sequence alignment (S116^2.67^Y, A233^5.52^S, Y277^6.44^F, W281^6.48^Y and K312^7.43^A). The mutated cone opsins were purified, and 11-*cis* retinal was added to reconstitute cone pigments in vitro. Interestingly, we observed a blue shift in the maximum absorption wavelength of the red-pigment upon residue substitution, bringing it closer to that of green-pigment (Figure 6D). It is worth noting that substitutions W281^6.48^Y and K312^7.43^A resulted in the absence of absorption wavelengths resembling those of either red-pigment or green-pigment (Figure 6D), elucidating the importance of the residues W281^6.48^ and K312^7.43^ for the spectral properties of red-pigment. These findings suggest that individual residue mutations may alter the photosensitivity of visual pigments.

## Discussion

Although rhodopsin has been extensively studied, there is a shortage of research on the human cone opsins, and the structures of cone opsins have not been elucidated yet. Moreover, the molecular mechanism of human color discrimination remains elusive. In this study, we solved the cryo-EM structures of three human cone opsins, namely LWS-opsin, MWS-opsin, and SWS-opsin, in complex with all-*trans* retinal and G_i_ proteins and reported the structural characteristics of human cone pigments. To our knowledge, this is the first report of the atomic structure of human cone opsins as well as cone pigments.

As an agonist, all-*trans* retinal interacts with cone opsins via a covalent bond and multiple hydrophobic interactions, inducing conformational changes of the “toggle switch” W^6.48^/Y^6.48^ of cone opsins and rearrangement of the conserved motifs in class A GPCRs. The R^3.50^ in the ERY motif stretches to TM5 and forms a hydrogen bond interaction with Y^5.58^ in active opsins. The outward shift of TM6 on the cytoplasmic side, facilitating its accommodation of G protein. The structural information revealed in our study is similar with that in rhodopsin-G_t_ complex (PDB ID 6OYA).

Based on structural analysis and mutagenesis experiments, we have elucidated the mechanism underlying the spectral tuning of the red-pigment. Specifically, mutations Y277^6.44^F, A233^5.52^S, and S116^2.67^Y in LWS-opsin resulted in a blue shift in the absorption spectrum of the red-pigment. In the LWS-opsin-G_i_ structure, Y277^6.44^ and C228^5.47^ in the retinal binding pocket form a hydrogen bond. The Y277^6.44^F mutation disrupts this interaction, potentially causing changes in retinal configuration, ultimately leading to a blue shift in the absorption spectrum. Similarly, the A233^5.52^S and S116^2.67^Y mutations both lead to absorption spectra shifts by influencing the interaction network of the retinal binding pocket. Of note, the substitution W281^6.48^Y may render the opsin incapable of normal activation, resulting in the loss of the UV absorption spectrum. As anticipated, the mutation of K312, which covalently binds to retinal, to alanine resulted in the absence of the absorption spectrum, as retinal could no longer bind to the opsin. These structural insights provide valuable information on the impact of mutations on the interaction between opsins and retinal.

Previous studies have reported that missense mutations in cone opsins associated with color vision deficiencies, however, the specific mechanisms remain unclear. (Weitz et al., 1992; Nathans et al., 1993; Ueyama et al., 2002; Carroll et al., 2004; Ueyama et al., 2004; Neitz et al., 2004; Mizrahi-Meissonnier et al., 2010; Cai et al., 2019; Neitz et al., 2020; Iarossi et al., 2021; Zhu et al., 2021). Our findings also provide some molecular insights from the structural perspective to color vision deficiencies. The most prevalent mutation in LWS-/MWS-opsin associated with red-green color blindness is C203^ECL2^R, disrupting the disulfide bond linkage between C203^ECL2^ and C126^3.25^. Notably, in both LWS-opsin and MWS-opsin, the residues W177^4.50^, N94^2.45^, and S143^3.42^ collaboratively form a hydrogen bonding network, and the mutations N94^2.45^K, S143^3.42^P, W177^4.50^R in LWS-opsin have been reported to induce protan color vision deficiency, whereas the mutations N94^2.45^K, W177^4.50^R in MWS-opsin result in deutan. Additionally, the mutations L56^1.54^P, G79^2.49^R, T190^ECL2^I, S214^5.52^P, P264^6.50^S, and R283^7.33^Q in SWS-opsin can lead to tritan color vision deficiency. These mutations may induce tritan color vision deficiency by causing significant steric hindrance or influencing the ligand-binding pocket. These analyses may provide insights into color vision deficiencies induced by mutations in cone opsins.

In summary, our study represents the first elucidation of the structures of three human cone opsins, enhancing our comprehension of cone pigments crucial for color perception. Subsequently, extensive mutagenesis studies will be undertaken to explore the influence of residue mutations on the stability of cone opsins and their UV-visible absorption characteristics.

## STAR ★ Methods

### Method details

#### Construct design and Protein Expression and purification

The human OPN1LW (UniProtKB-P04000)/OPN1MW (UniProtKB-P04001)/OPN1SW (UniProtKB-P03999) genes was cloned into a pFastBac vector using ClonExpress II One Step Cloning Kit (Vazyme). The vector is characterized by an N-terminal hemagglutinin (HA) signal sequence and a Flag tag (DYKDDDD), while the C-terminus is comprised of a tobacco etch virus (TEV) protease cleavage site, followed by a 10×His tag. To improve protein yield and homogeneity, a thermostabilized apocytochrome b562 (BRIL)(Chun et al., 2012) was inserted before the OPN1LW/OPN1MW/OPN1SW, and a constitutive active mutation E129^3.28^Q was introduced (constitutive active mutation E110Q^3.28^ in OPN1SW)(Zhukovsky and Oprian, 1989; Sakmar et al., 1989; Robinson et al., 1992; Standfuss et al., 2011). For cryo-EM studies, a NanoBiT tethering strategy was employed for complex stabilization(Duan et al., 2020; Zhou et al., 2020). The sequence of human OPN1LW/OPN1MW/OPN1SW was fused with a LgBiT subunit (Promega) at the C-terminus. A dominant-negative Gα_i1_ (DNGα_i1_) was generated by introducing five mutations, S47C, G202T, G203A, E245A, and A326S to decrease the affinity of nucleotide-binding(Liu et al., 2016; Draper-Joyce et al., 2018). Human Gβ_1_ was fused with an N-terminal 6× His tag and a C-terminal HiBiT, while bovine Gγ was employed. The DNGα_i1_ and Gβ_1_γ_2_ subunits were cloned into pFastbac1 and pFastBac Dual vectors, respectively. Additionally, the scFv16 gene was cloned into the pFastBac1 vector, incorporating an N-terminal GP67 signaling peptide and a C-terminal HRV3C protease cleavage site followed by an 8× His-tag(Koehl et al., 2018; Mao et al., 2020).

The modified OPN1LW/OPN1MW/OPN1SW was co-expressed with DNGα_i1_, Gβ_1_γ_2_ and scFv16 in Sf9 insect cells (Invitrogen) using Bac-to-Bac Baculovirus Expression System (Invitrogen). The cells at a density of 1.5×10^6^ cells/mL were infected with high-titre viral stocks (>10^9^ viral particles/mL) at a multiplicity of infection (MOI) ratio of 1:1:1:1 for OPN1LW/OPN1MW/OPN1SW, DNGα_i1_, Gβ_1_γ_2_ and scFv16. Cells were grown at 27 °C for 48 h and then harvested by centrifugation. The cell pellets were stored at –80 °C until use.

The cells expressing the cone opsin-G_i_ complex were thawed on ice and suspended in a buffer containing 20 mM HEPES, pH 7.5, 100 mM NaCl, 10 mM MgCl_2_, 20 mM KCl, 5 mM CaCl_2_, 100 μM TCEP, 5 μM all-*trans* retinal, and EDTA-free protease inhibitor cocktail tablets (Roche) by dounce homogenization. Then, the suspension was incubated at room temperature for 1 h with apyrase (25 mU/mL; New England Biolabs). The supernatant was removed by ultracentrifugation at 32,000 rpm for 30 min, and the membrane pellet was resuspended in a buffer containing 20 mM HEPES, pH7.5, 100 mM NaCl, 5 mM CaCl_2_, 100 μM TCEP, 5% glycerol, 5 μM all-*trans* retinal, EDTA-free protease inhibitor and apyrase (50 mU/mL). The membranes were then solubilized with the addition of 0.5% (w/v) lauryl maltose neopentyl glycol (LMNG, Anatrace) and 0.05% (w/v) cholesteryl hemisuccinate (CHS, Anatrace). After gentle stirring for 2 h at 4 °C, the supernatant was isolated by ultracentrifugation at 32,000 rpm for 40 min, and then incubated with TALON IMAC resin (Clontech) at 4 °C for 2 h in the presence of 20 mM imidazole. The resin was washed with 10 column volumes of Wash Buffer I containing 20 mM HEPES, pH 7.5, 100 mM NaCl, 5% glycerol, 0.1% LMNG, 0.01% CHS, 100 μM TCEP, 20 mM imidazole, and 5 μM all-*trans* retinal. Followed by 10 column volumes of Wash Buffer II containing 20 mM HEPES, pH 7.5, 100 mM NaCl, 5% glycerol, 0.05% LMNG, 0.005% CHS, 100 μM TCEP, 30 mM imidazole, and 5 μM all-*trans* retinal. The complex was then eluted by 3 column volumes of Elution Buffer containing 20 mM HEPES, pH 7.5, 100 mM NaCl, 5% glycerol, 0.03% LMNG, 0.003% CHS, 100 μM TCEP, 300 mM imidazole, and 5 μM all-*trans* retinal. The protein was concentrated and subjected onto a Superose 6 Increase 10/300 column (GE Healthcare) in the buffer containing 20 mM HEPES, pH 7.5, 100 mM NaCl, 0.003% LMNG, 0.0003% CHS, 100 μM TCEP, 5 μM all-*trans* retinal. The purified complex fractions were collected and concentrated using a 100 kDa molecular mass cut-off concentrator (Millipore) for cryo-EM grid preparation.

### Cryo-EM sample preparation and data acquisition

To prepare the cryo-EM grid of the LWS-opsin-G_i_ complex, 3.5 μL of the purified samples at 10 mg/mL were added to 300 Mesh R1.2/1.3 Cu Quantifoil grids (glow discharged at 15mA for 40 seconds with a Glow discharge cleaning system). Grid were blotted with qualitative filter paper in a Vitrobot Mark Ⅳ (Thermo Fisher Scientific) at 4℃ and 100% humidity for 3 second using a blot force of –2 prior to plunging into liquid ethane. For cryo-EM data acquisition, grids were loaded on a Thermo Fisher Scientific 300 kV TEM Titan Krios equipped with Gatan K3 direct electron detector. Raw movies were used to record at a magnification of 130,000 and the pixel size was 0.334 Å. The movie stacks were automatically acquired with the defocus range from –1.8 to –2.2 μm and collected using SerialEM in super-resolution mode. The exposure time was 2.5 s, with frames collected for a total of 50 frames (0.05 s/frame) per sample with a total dose of 41 e−/Å^2^.

For MWS-opsin-G_i_ complex, 3.0 μL of the purified samples at 7 mg/mL were added to 300 Mesh R1.2/1.3 Au Quantifoil grids. For SWS-opsin-G_i_ complex, 3.5 μL of the purified samples at 8 mg/mL were added to 300 Mesh R1.2/1.3 Cu Quantifoil grids. All grids were blotted with qualitative filter paper in a Vitrobot Mark (Thermo Fisher Scientific) at 4℃ and 100% humidity for 3 seconds and plunge frozen in liquid ethane, and then stored in liquid nitrogen until checked. We used the 300 kV Titan Krios Gi3 microscope (Thermo Fisher Scientific FEI, the Kobilka Cryo-EM Center of the Chinese University of Hong Kong, Shenzhen) to check the grids and collect the cryo-EM data of the all complexes. The Gatan K3 BioQuantum camera at a magnification of 105,000 was used to record movies, and the pixel size was 0.83 Å. The movie stacks were automatically acquired with the defocus range from –1.0 to –2.0 μm. The exposure time was 2.5 s, with frames collected for a total of 50 frames (0.05 s/frame) per sample. For the MWS-opsin-G_i_ complex, the total dose is 54 e−/Å^2^ and for the SWS-opsin-G_i_ complex, the total dose is 51.9 e−/Å^2^. SerialEM 3.7 was used for semiautomatic data acquisition.

### Cryo-EM data processing

For the LWS-opsin-G_i_ complex, a total of 5,292 movies stacks were imported into cryoSPARC V3.3.2. After motion corrected, electron-dose weighted and CTF estimation filtered out movies with a resolution greater than 4 Å. The initial particle was performed by cryoSPARC blob picker. A total number of 1,550,865 particles were extracted with the blob picking from 500 random movies. After two rounds of 2D classification, the good 2D classes were used for template picker. A total number of 4,521,699 particles were extracted with the template picking from all movies. After four rounds of 2D classification, the good 2D particles were reduce to 1,244,445. The number of particles was further reduced to 241,247 by three rounds heterogeneous refinement and Ab-initio reconstruction. The nonuniform refinement enables us to reconstruct the 3.41 Å structure. To further improve the resolution, the final particle sets were re-extracted with the original box size and further applied for final nonuniform refinement and local refinement in cryoSPARC. A density map was obtained with an overall resolution of 3.35 Å (determined by gold standard Fourier shell correlation (FSC) using the 0.143 criterion).

For the MWS-opsin-G_i_ complex, a total of 5,166 movies stacks were imported into cryoSPARC V4.4.1. After motion corrected, electron-dose weighted and CTF estimation filtered out movies with a resolution greater than 4 Å. The initial particle was performed by cryoSPARC blob picker. A total number of 6,240,737 particles were extracted with the blob picking. After four rounds of 2D classification, the number of good particles was reduced to 1,443,488. The number of particles was further reduced to 579,445 by heterogeneous refinement and Ab-initio reconstruction. The nonuniform refinement enables us to reconstruct the 3.43 Å structure. To further improve the resolution, the final particle sets were re-extracted with the original box size and further applied for final nonuniform refinement and local refinement in cryoSPARC. A density map was obtained with an overall resolution of 2.48 Å (determined by gold standard Fourier shell correlation (FSC) using the 0.143 criterion).

For the SWS-opsin-G_i_ complex, a total of 6,084 movies stacks were imported into cryoSPARC V4.4.1. After motion corrected, electron-dose weighted and CTF estimation filtered out movies with a resolution greater than 4 Å. The initial particle was performed by cryoSPARC blob picker. A total number of 3,841,818 particles were extracted with the blob picking. After four rounds of 2D classification, the number of good particles was reduced to 1,833,322. The number of particles was further reduced to 407,822 by heterogeneous refinement and Ab-initio reconstruction. The nonuniform refinement enables us to reconstruct the 3.43 Å structure. To further improve the resolution, the final particle sets were re-extracted with the original box size and further applied for final nonuniform refinement and local refinement in cryoSPARC. A density map was obtained with an overall resolution of 2.61 Å (determined by gold standard Fourier shell correlation (FSC) using the 0.143 criterion).

### Model building and refinement

For the complexes, the initial model for the cone opsins was the Cryo-EM structure of Rhodopsin (PDB ID 6CMO)(Kang et al., 2018) and the initial model for the G_i_-scFv16 complex was generated from the CCR1-G_i_ complex (PDB ID 7VL8)(Shao et al., 2022). The models were docked into the cryoEM density map using UCSF Chimera(Pettersen et al., 2004), followed by iterative manual adjustment and rebuilding in COOT(Emsley and Cowtan, 2004) according to side-chain densities. Real-space refinement was performed using Phenix programs(Adams et al., 2010). The model statistics were validated using MolProbity(Chen et al., 2010). Structural figures were prepared in Chimera and PyMOL (https://pymol.org/2/). The final refinement statistics are provided in Supplementary Table S1.

### Expression of cone opsins in Sf9 cells and pigments reconstitution

Expression constructs of human cone opsins (WT and mutants) were inserted into the previously described pFastBac vector. Following this, human cone opsins were expressed in Sf9 cells utilizing the Bac-to-Bac baculovirus expression system. After 48h of expression, the infected Sf9 cells were harvested and stored at –80 °C until use.

To isolate membranes, frozen Sf9 cells were first thawed and disrupted in a Dounce homogenizer and a hypotonic buffer composed of 10 mM HEPES, pH 7.5, 10 mM MgCl_2_, 20 mM KCl and EDTA-free protease inhibitor. Membranes then were pelleted by centrifugation at 32,000 rpm for 30 min. Washed membranes were resuspended and solubilized in a buffer containing 50 mM HEPES, pH 7.5, 150 mM NaCl, 10% glycerol, 0.5% LMNG, 0.05% CHS and EDTA-free protease inhibitor. The

suspension was rotated for 2 h, and insoluble material was removed by centrifugation at 32,000 rpm for 40 min. The supernatant was incubated with TALON IMAC resin (Clontech) at 4 °C for 2 h in the presence of 20 mM imidazole. The resin was packed into a glass column and washed with 10 column volumes of Wash Buffer A (50 mM HEPES, pH 7.5, 150 mM NaCl, 10% glycerol, 0.05% LMNG, 0.005% CHS and 20 mM imidazole), followed by 10 column volumes of Wash Buffer B (50 mM HEPES, pH 7.5, 150 mM NaCl, 10% glycerol, 0.05% LMNG, 0.005% CHS and 30 mM imidazole), and was then eluted with 3-5 column volumes of the Elution buffer (50 mM HEPES, pH 7.5, 150 mM NaCl, 10% glycerol, 0.05% LMNG, 0.005% CHS and 300 mM imidazole).

Cone pigments were reconstituted with purified cone opsins and 11-*cis* retinal. Retinal was added to the cone opsin at molar ratio of 1:1 and then incubated in the dark for 1 h at 4 °C.

### UV-visible spectroscopy of cone pigments

UV-visible spectra of freshly purified cone pigment samples (WT and mutants) were recorded in the dark with a QuickDrop UV-visible spectrophotometer (Molecular Devices). To validate the covalent Schiff base between cone opsins and retinal, UV-visible spectra of protein samples involving LWS-opsin-G_i_, MWS-opsin-G_i_, and SWS-opsin-G_i_, in the absence or presence of acid (acidified to pH 2), were measured with a flexstation3 (Molecular Devices). The difference spectra of purified cone pigments-G_i_ was derived by subtracting the spectra recorded before acidification from the spectra recorded after acidification.

## Acknowledgments

J.Z. was supported by the National Natural Science Foundation of China (grant no. 32271260), the CAS ‘‘Light of West China’’ Program (xbzg-zdsys-202005), and the Natural Science Foundation of Jiangxi Province (grant no. 20224ACB206046). Y.L. is an investigator of SUSTech Institute for Biological Electron Microscopy. Y.L. was supported by the National Natural Science Foundation of China (grant no.32322004; grant no. 32171205; grant no. U22A20338), and the Shenzhen Natural Science Foundation (grant no. 20220815130429001).

## Author contributions

Q.P. performed the experiments (including designing and generating all recombinant DNA plasmids, optimizing expression and purification of cone opsin-G_i_ complexes, determining the UV-Visible absorption spectra), interpreted the data, prepared the figures, and wrote the first draft of the manuscript (ms) J.L. contributed to building the atomic model on the basis of cryo-EM maps and revised the ms; H.J. contributed to the revision of the ms; X.C. carried out cryo-EM data processing; Q.L. contributed to cryo-EM experiments. S.Z. prepared mutants; Y.Z contributed to cryo-EM data acquisition; W.N. contributed to the expression of Sf9 cells and the harvesting cells; J.Z. initiated the project, conceived the study, designed the experiments, interpreted the data, wrote the ms and supervised for the research. The other authors all coordinated the study.

## Supplementary information

**Figure S1.**
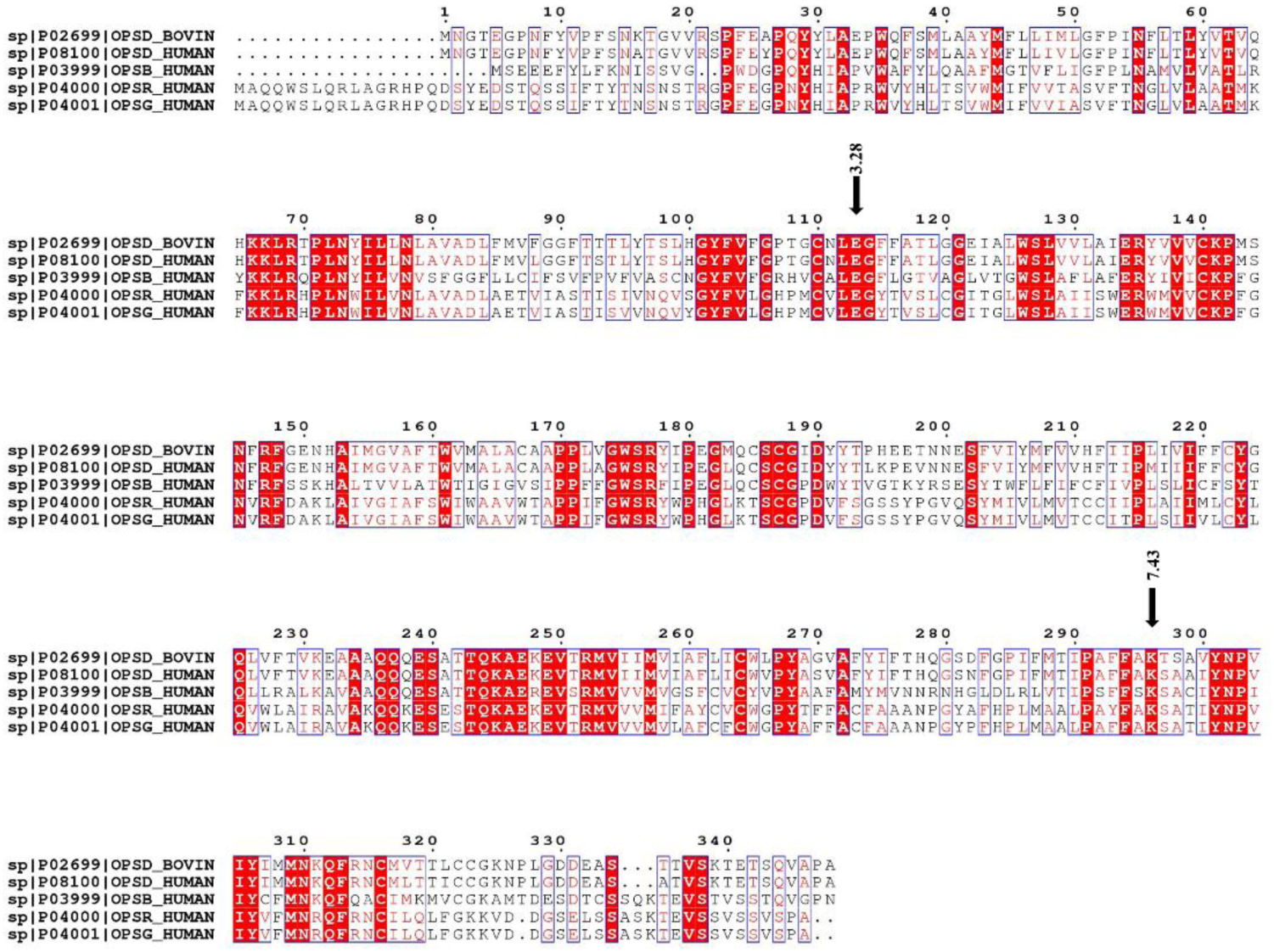
Sequence alignment of Rhodopsin with OPN1LW, OPN1MW and OPN1SW. The figure presents the result of the sequence alignment of Rhodopsin and OPN1LW, OPN1MW and OPN1SW (also known as OPSD, OPSR, OPSG, and OPSB respectively). The black arrows highlight the conserved E^3.28^ and K^7.43^ amino acid residues among bovine/human rhodopsin and the three human cone opsins.

**Figure S2.**
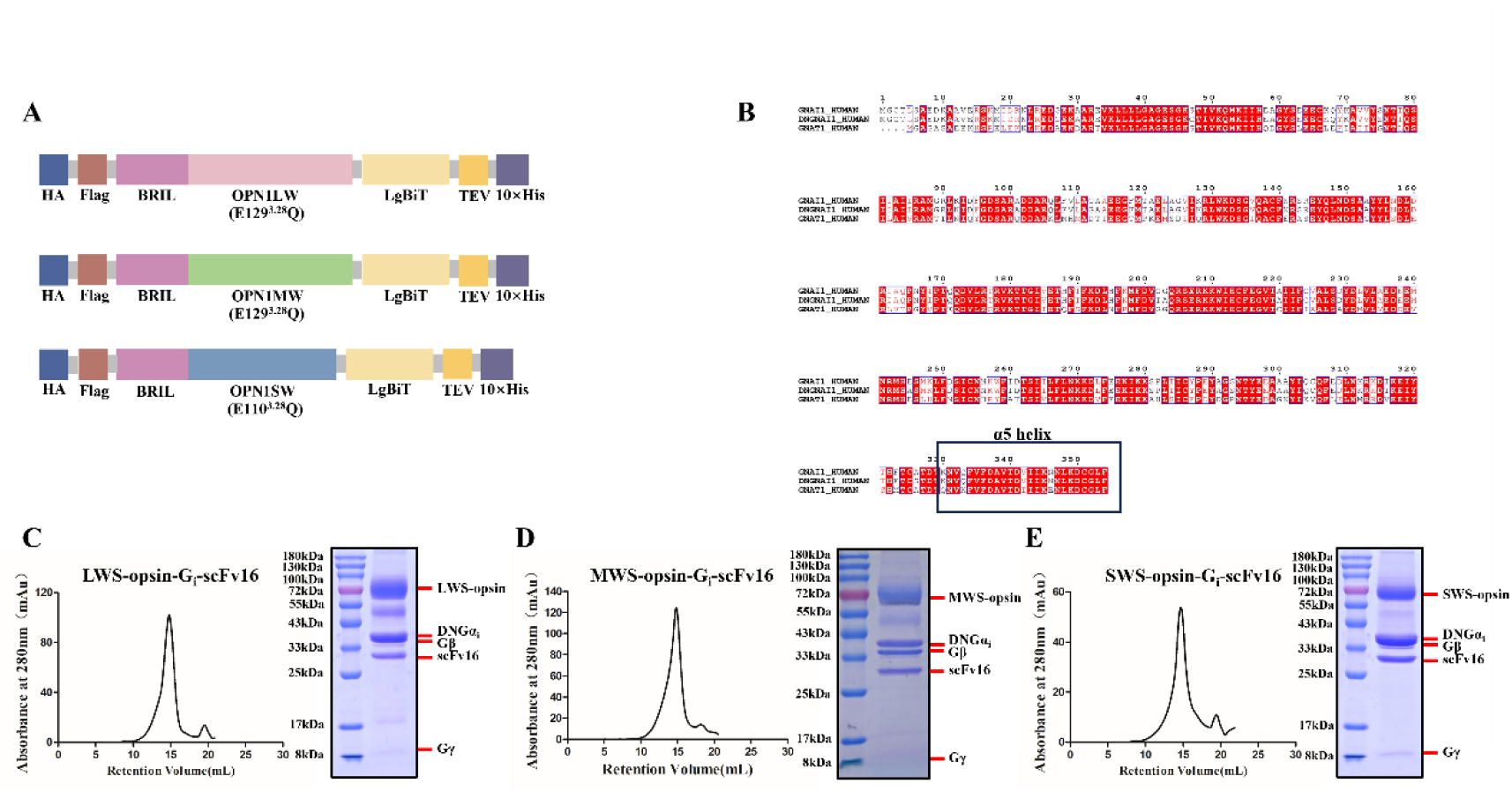
Construct design and purification of LWS-opsin-G_i_, MWS-opsin-G_i_ and SWS-opsin-G_i_ complexes. (**A**) Construct design of OPN1LW, OPN1MW and OPN1SW. **(B)** Sequence alignment of DNGα_i1_ with Gα_i1_ and Gα_t1_. **(C)** Representative elution profile and SDS-PAGE analysis of the LWS-opsin-G_i_ complex. **(D)** Representative elution profile and SDS-PAGE analysis of the MWS-opsin-G_i_ complex. **(E)** Representative elution profile and SDS-PAGE analysis of the SWS-opsin-G_i_ complex.

**Figure S3.**
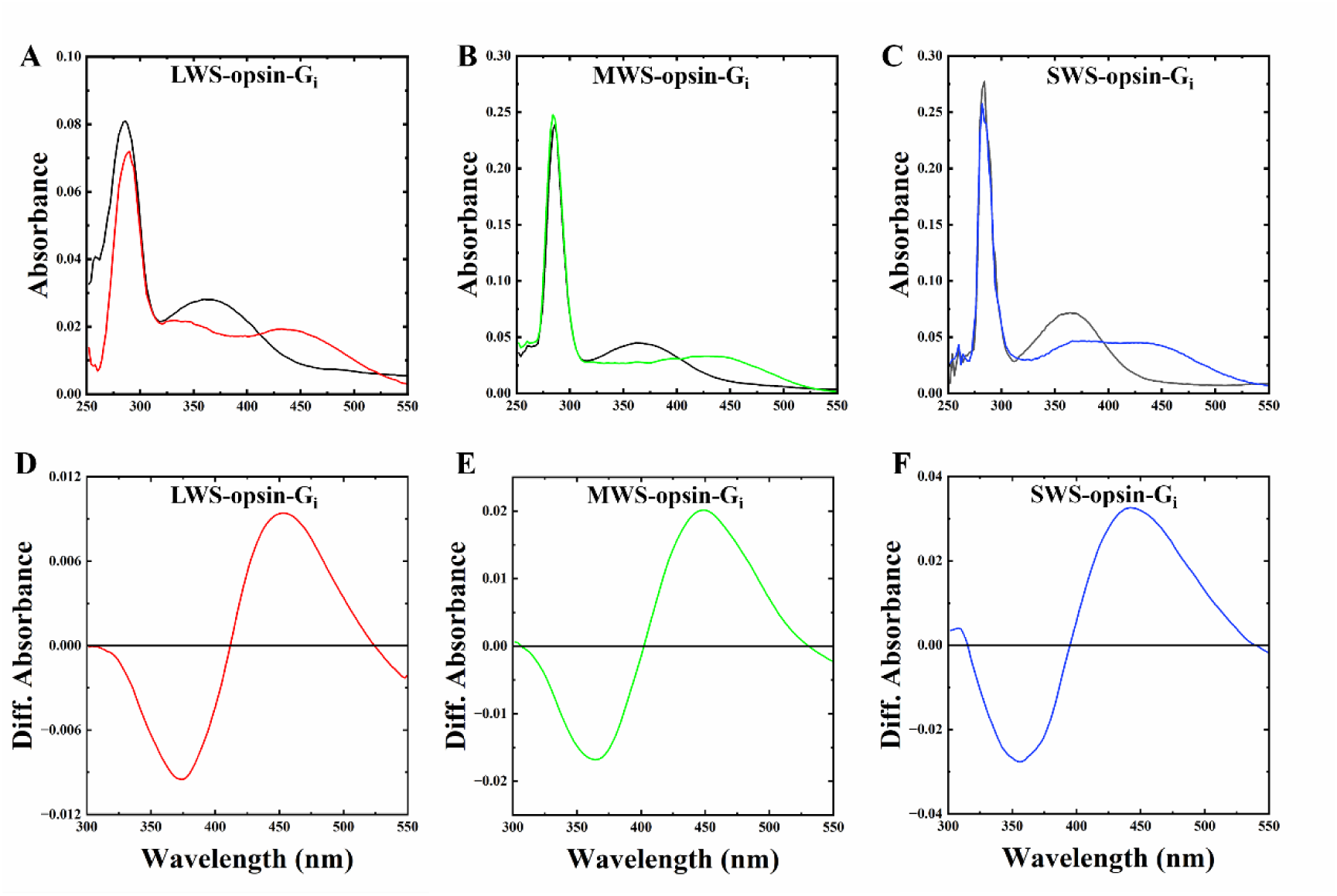
UV-visible spectra and difference spectra of LWS-opsin-G_i_, MWS-opsin-G_i_, and SWS-opsin-G_i_. (**A-C**) UV-visible spectra of LWS-opsin-G_i_, MWS-opsin-G_i_, and SWS-opsin-G_i_ complexes were measured. Spectra in the absence (black curve) or presence of acid (pH 2.0). **(D-F)** Difference spectra of purified cone pigments-G_i_ were derived by subtracting pre-acidification recorded spectra from those recorded post-acidification. Generation of a product absorbing at 440 nm indicates a protonated Schiff base and thus a covalent linkage between retinal and lysine (red curve for LWS-opsin-G_i_, green curve for MWS-opsin-G_i_ and blue curve for SWS-opsin-G_i_).

**Figure S4.**
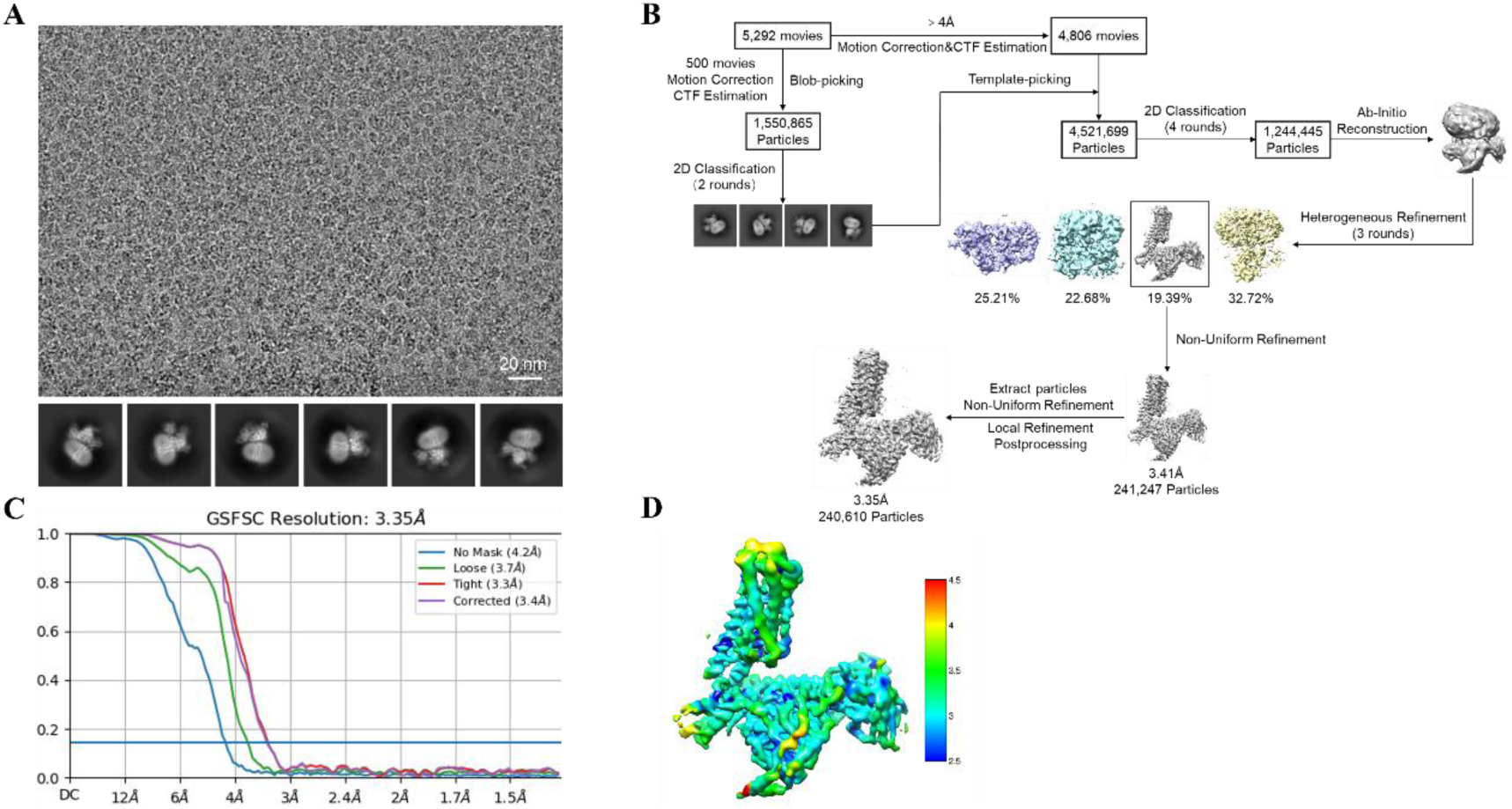
Single particle cryo-EM analysis of the LWS-opsin-G_i_ complex. (**A**) Representative cryo-EM micrograph of LWS-opsin-G_i_ complex and the selected two-dimensional class averages of cryo-EM particle images. **(B)** Cryo-EM workflow chart of data processing. **(C)** Gold-standard Fourier shell correlation (FSC) curve showing an overall resolution at 3.4 Å for the LWS-opsin-G_i_ complex. **(D)** The density map of LWS-opsin-G_i_ colored by local resolution (Å).

**Figure S5.**
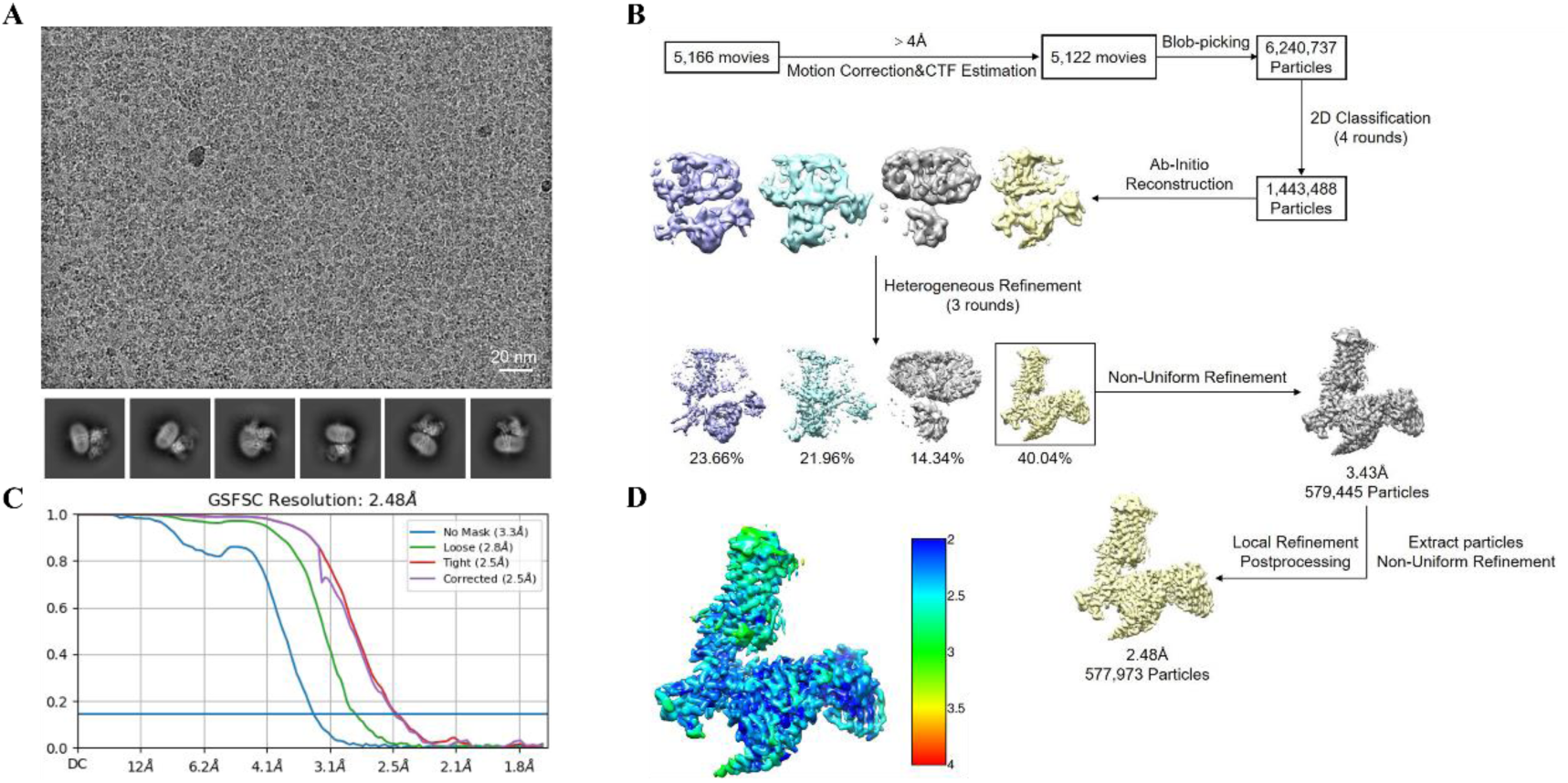
Single particle cryo-EM analysis of the MWS-opsin-G_i_ complex. (**A**) Representative cryo-EM micrograph of MWS-opsin-G_i_ complex and the selected two-dimensional class averages of cryo-EM particle images. **(B)** Cryo-EM workflow chart of data processing. **(C)** Gold-standard Fourier shell correlation (FSC) curve showing an overall resolution at 2.5 Å for the MWS-opsin-G_i_ complex. **(D)** The density map of MWS-opsin-G_i_ colored by local resolution (Å).

**Figure S6.**
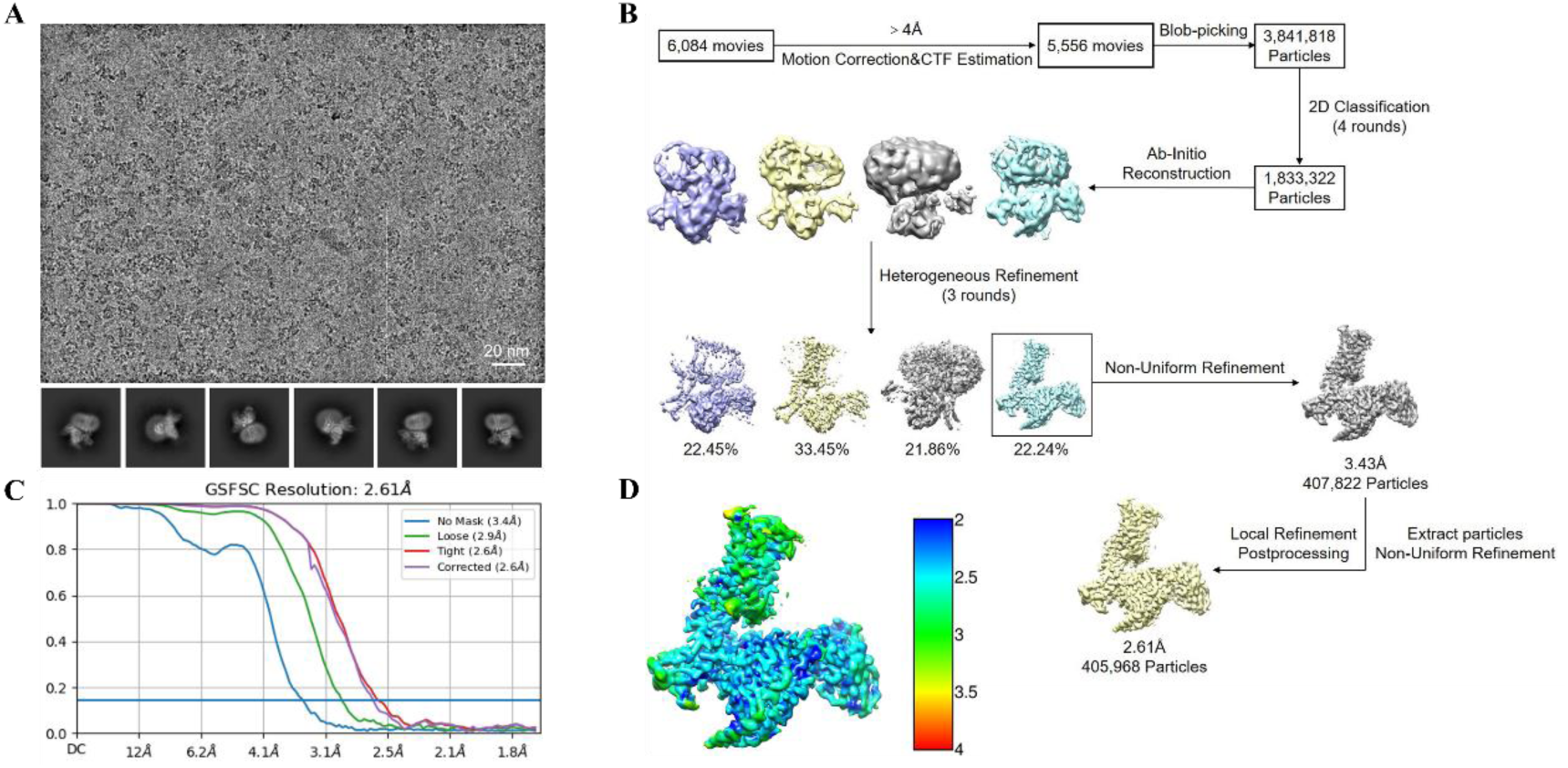
Single particle cryo-EM analysis of the SWS-opsin-G_i_ complex. (**A**) Representative cryo-EM micrograph of SWS-opsin-G_i_ complex and the selected two-dimensional class averages of cryo-EM particle images. **(B)** Cryo-EM workflow chart of data processing. **(C)** Gold-standard Fourier shell correlation (FSC) curve showing an overall resolution at 2.6 Å for the SWS-opsin-G_i_ complex. **(D)** The density map of SWS-opsin-G_i_ colored by local resolution (Å).

**Figure S7.**
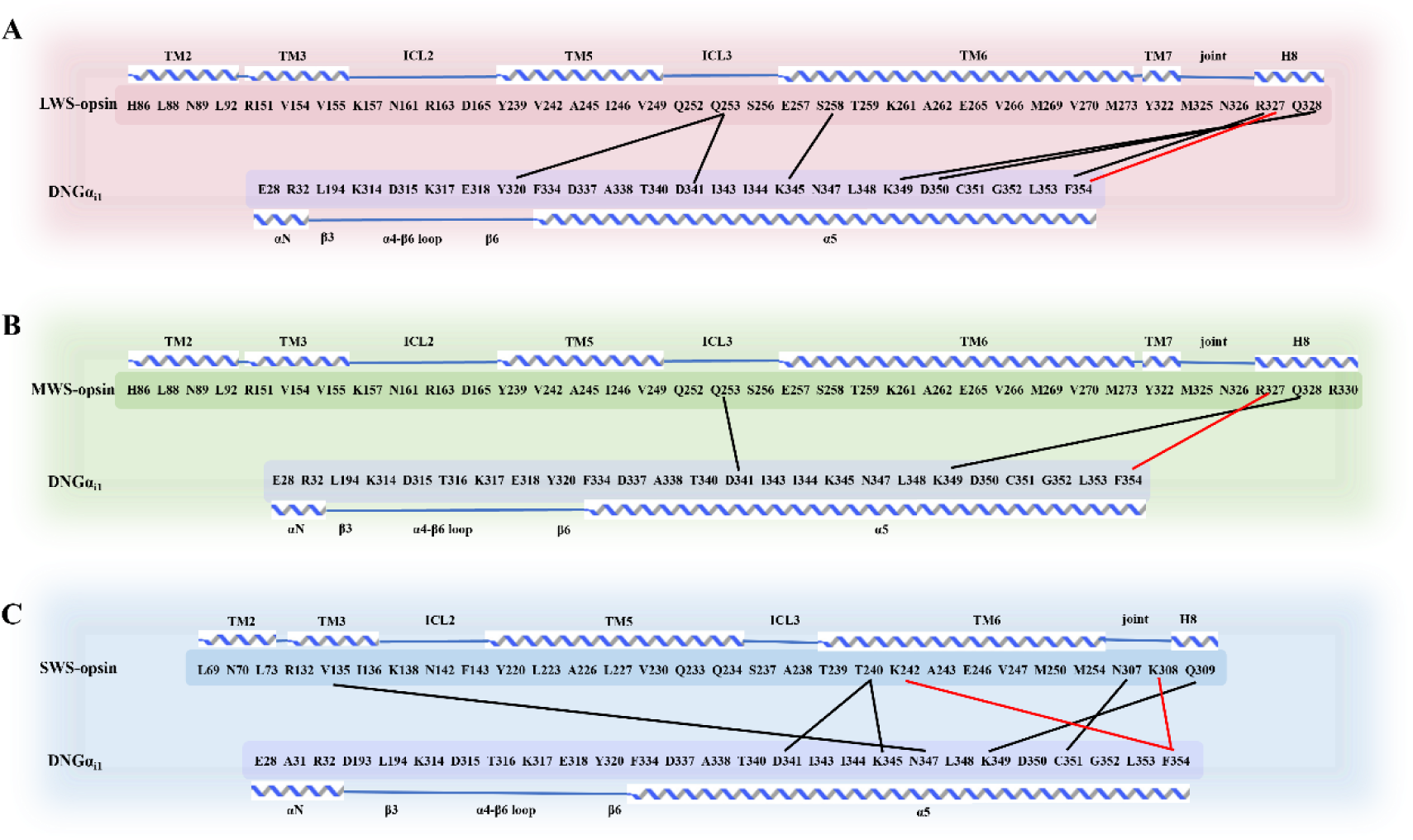
Interface between LWS-opsin/MWS-opsin/SWS-opsin and G_i_. (**A-C**) The connective interaction map illustrating the receptor/Gα_i_ interaction in the LWS-opsin-G_i_ complex **(A)**, MWS-opsin-G_i_ complex **(B)** and SWS-opsin-G_i_ complex **(C)**. The black line marks polar interactions and the red line marks π-cation interaction.

**Figure S8.**
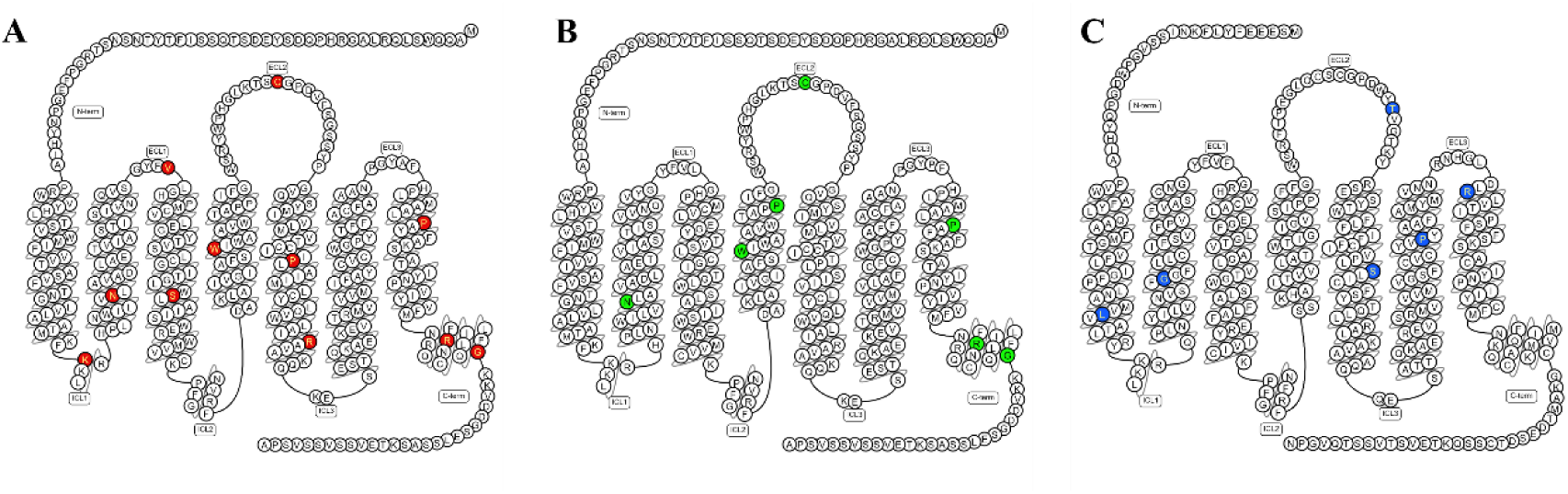
Color Vision Deficiency Mutations in Cone Opsins. (**A-C**) Snake plots of LWS-opsin **(A)**, MWS-opsin **(B)**, and SWS-opsin **(C)**, with highlighted residues indicating mutations associated with color vision deficiencies.

**Table S1.** Cryo-EM data collection, refinement and validation statistics.

|  | LWS-opsin-G<sub>i</sub><br>(EMD-35714)<br>(PDB 8IU2) | MWS-opsin-G<sub>i</sub><br>(EMD-38800)<br>(PDB 8Y01) | SWS-opsin-G<sub>i</sub><br>(EMD-38801)<br>(PDB 8Y02) |
| --- | --- | --- | --- |
| **Data collection and processing** |  |  |  |
| Magnification | 130,000 | 105,000 | 105,000 |
| Voltage (kV) | 300 | 300 | 300 |
| Electron exposure (e-/Å<sup>2</sup>) | 41 | 54 | 51.9 |
| Defocus range (μm) | -1.8--2.2 | -1.0--2.0 | -1.0--2.0 |
| Pixel size (Å) | 0.668 | 0.83 | 0.83 |
| Symmetry imposed | *C1* | *C1* | *C1* |
| Initial particle images (#) | 4,521,699 | 6,240,737 | 3,841,818 |
| Final particle images (#) | 240,610 | 577,973 | 405,968 |
| Map resolution (Å) | 3.35 | 2.48 | 2.61 |
| FSC threshold | 0.143 | 0.143 | 0.143 |
| **Refinement** |  |  |  |
| Initial model used (PDB code) | 6CMO, 7VL8 | 6CMO, 7VL8 | 6CMO, 7VL8 |
| Model resolution (Å) | 4.5 | 4.5 | 4.5 |
| FSC threshold | 0.143 | 0.143 | 0.143 |
| Model composition |  |  |  |
| Non-hydrogen atoms | 8737 | 8740 | 8385 |
| Protein residues | 1137 | 1137 | 1093 |
| Ligands | 1 | 1 | 1 |
| r.m.s. deviations |  |  |  |
| Bond lengths (Å) | 0.01 | 0.01 | 0.01 |
| Bond angles (°) | 1.34 | 1.30 | 0.97 |
| Validation |  |  |  |
| MolProbity score | 1.56 | 1.54 | 1.57 |
| Ramachandran plot |  |  |  |
| Favored (%) | 95.09 | 95.63 | 95.26 |
| Allowed (%) | 4.91 | 4.37 | 4.73 |
| Disallowed (%) | 0.00 | 0.00 | 0.00 |

## Notes

### Competing Interest Statement

The authors have declared no competing interest.

